# Histone H2BK120 ubiquitination modulates PRC1 activity and H2AK119ub deposition on nucleosomes

**DOI:** 10.64898/2026.02.20.706939

**Authors:** Ying Yu, Dan Cai, Yuzhu Zhang, Guo Bai, Xinyi Wan, Yu Liu, Fan Wang, Zhuowei Tian, Jianglai Nie, Juan Chen, Han Xue, Ming Lei, Zhiyuan Zhang, Chi Yang, Dong Fang, Jing Huang

## Abstract

Polycomb group and Trithorax group complexes antagonistically regulate developmental gene expression, yet the mechanisms coordinating their interplay remain poorly understood. Here, we uncover a synergistic histone crosstalk in which H2BK120ub (histone H2B ubiquitination at lysine 120), a Trithorax-associated active mark, enhances Polycomb repressive complex 1 (PRC1) activity by directly binding specific PCGF subunits of PRC1 on the nucleosome surface, as shown in cryo-electron microscopy (cryo-EM) analysis. Beyond regulating enzymatic activity, H2BK120ub also modulates the competitive binding between the catalytic RNF2-BMI1 module and the RYBP subunit of noncanonical PRC1.4 complexes to nucleosomes, potentially restricting H2AK119ub (histone H2A ubiquitination at lysine 119) spreading to neighboring nucleosomes. In mouse embryonic stem cells, H2BK120ub directs the deposition of H2AK119ub within the gene bodies of a subset of developmental genes, which exhibit more rapid activation compared to canonical H3K4me3/H3K27me3 bivalent genes. These findings reveal a previously unrecognized mechanism that links active and repressive histone modifications, providing new insights into the intricate chromatin regulation of gene expression during development.

## Introduction

Cell fate determination and maintenance during development rely on precise gene expression programs orchestrated by chromatin modulators. Among these, the evolutionarily conserved Polycomb group (PcG) and Trithorax group (TrxG) proteins emerge as key players with antagonistic functions in transcriptional regulation^1–5^. PcG proteins assemble into Polycomb repressive complex 1 (PRC1) and Polycomb repressive complex 2 (PRC2) to catalyze the ubiquitination of histone H2A on lysine 119 (H2AK119ub) and the methylation of histone H3 on lysine 27 (H3K27me), respectively^6–9^. These complexes employ multiple positive feedback loops to reinforce their individual and synergistic histone modification activities, facilitating the establishment and maintenance of repressive transcriptional states^10–17^. On the contrary, TrxG proteins, including the SWI/SNF chromatin remodeling complex and the histone methyltransferase MLL/COMPASS family, counteract the repressive effects of the PcG complexes and promote gene activation^18–20^. The MLL/COMPASS-catalyzed methylation of histone H3 on lysine 4 (H3K4me) is stimulated by the active histone mark H2BK120ub (ubiquitination of histone H2B at lysine 120), coordinating gene activation regulation^21–24^. Beyond their individual opposing functions, PcG and TrxG complexes co-mark the promoters of developmental genes with both H3K27me3 and H3K4me3 modifications. This co-regulation maintains genes in a poised state, enabling rapid activation or repression during cell lineage determination and playing a crucial role in embryogenesis^25–28^.

Mammalian PcG machinery, particularly PRC1, has expanded into a variety of complexes mainly classified into canonical (cPRC1) and noncanonical (ncPRC1) variants. All PRC1 complexes contain a catalytic module composed of the E3 ligase RING1A/1B and one of the six PCGF proteins (PCGF1-PCGF6), both essential for catalyzing the repressive H2AK119ub modification^10,11^. In the cPRC1 complexes, the *Drosophila* Polycomb (Pc) protein has evolved into its mammalian Chromobox (CBX) orthologs, which specifically recognize PRC2-deposited H3K27me3 marks. This recognition, in concert with the Polyhomeotic-like proteins (PHC1-3), facilitates the formation of higher-order repressive chromatin domains, commonly referred to as Polycomb bodies^29–35^. In the ncPRC1 complexes, each of the six PCGF proteins assembles with distinct auxiliary subunits to generate ncPRC1.1 through ncPRC1.6 variants. These complexes incorporate RYBP or YAF2, which are mutually exclusive with the cPRC1-specific CBX subunit, to recognize H2AK119ub-modified nucleosomes and promote long-distance spreading of H2AK119ub marks, maintaining repressive chromatin domains^11,12,36^.

Previous studies have provided insights into the histone crosstalk mechanisms within PcG and TrxG complexes individually. However, the cooperative interplay between these complexes, particularly at development-related bivalent genes, remains poorly characterized. Here, we present cryo-electron microscopy (cryo-EM) structures of cPRC1 and ncPRC1 complexes bound to *in vitro* ubiquitinated nucleosomes. These structures provide mechanistic insight into how PRC1 bridges gene activation and repression by integrating the TrxG-associated active H2BK120ub mark to promote the deposition of the repressive H2AK119ub mark and to prevent the binding of the RYBP subunit and potentially fine-tune the propagation of H2AK119ub.

## Results

### H2BK120ub enhances H2AK119ub deposition via direct interaction with cPRC1 complex

To investigate the crosstalk between histone marks H2AK119ub and H2BK120ub, which play pivotal roles in Polycomb and Trithorax-mediated epigenetic control, we carried out *in vitro* ubiquitination assays to examine how the presence of one histone ubiquitination affects the deposition of the other on the nucleosome. Our results showed that pre-existing H2BK120ub notably promoted the deposition of H2AK119ub by the catalytic module of human cPRC1 complex. Conversely, the presence of H2AK119ub impaired the enzymatic activity of human RNF20-RNF40 complex, which serves as the E3 ligase responsible for the H2BK120ub modification (Fig. 1a-c).

**Figure 1.**
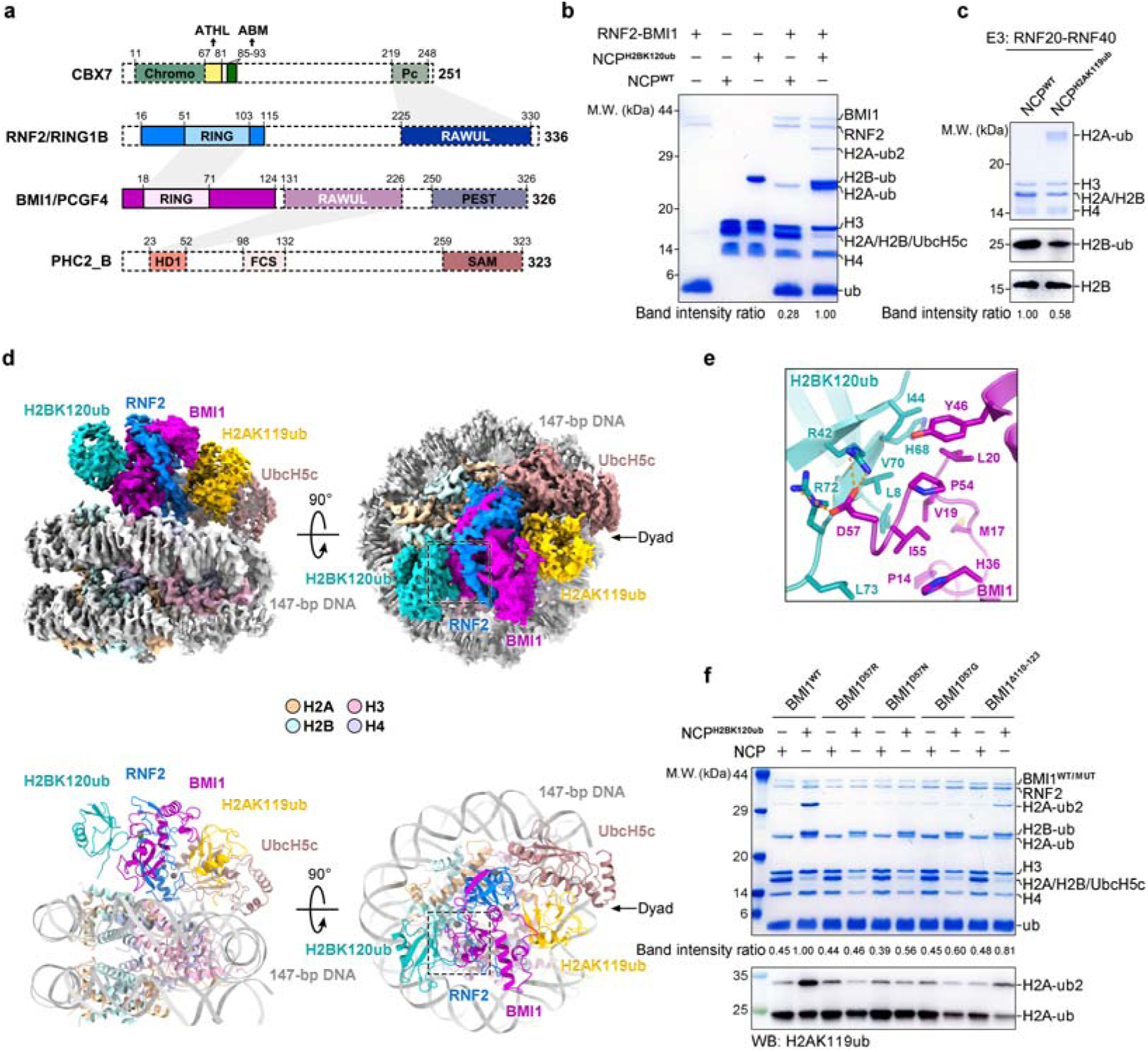
H2BK120ub interacts with cPRC1 complex, promoting the deposition of H2AK119ub on nucleosomes. **a**, Schematic of the domain organizations of the human cPRC1 complex. Abbreviations of domain names: Chromo, chromodomain; ATHL, AT-hook like; ABM, acidic-patch-binding motif; Pc, Polycomb box; RAWUL, ring finger and WD40 associated ubiquitin-like; PEST, proline, glutamic acid, serine and threonine rich; HD1, homology domain 1; FCS, zinc finger with a characteristic phenylalanine–cysteine–serine sequence motif; SAM, sterile alpha motif. **b**, *In vitro* ubiquitination assays of RNF2-BMI1 with unmodified or H2BK120ub-modified nucleosomes, visualized by Coomassie blue staining. The bottom row represents the ratio of the band intensity of H2A-ub (including H2A-ub and H2A-ub2) to that of H3, normalized to lane 5 (BMI1-NCP^H2BK120ub^, set to 1.00). The experiment was repeated at least three times with similar results **c**, *In vitro* ubiquitination assays of RNF20-RNF40 with unmodified or H2AK119ub-modified nucleosomes, analyzed by Western blot. The bottom row represents the ratio of the band intensity of H2B-ub to that of H3, normalized to lane 1 (RNF20-RNF40-NCP^H2AK119ub^, set to 1.00). The experiment was repeated at least three times with similar results **d**, Overall structure of cPRC1-E2∼ub-NCP^H2BK120ub^ complex. Cryo-EM density map (upper panel) and atomic model (lower panel) of cPRC1-E2∼ub-NCP^H2BK120ub^ complex are shown from two orthogonal views. The cryo-EM map is segmented according to the components of the cPRC1-E2∼ub-NCP^H2BK120ub^ complex. The color scheme of the cPRC1-E2∼ub-NCP^H2BK120ub^ complex is the same as depicted in (a), and the 147-bp DNA chains are shown in light and dark gray, respectively. Ub, ubiquitin; E2, UbcH5c. **e**, Detailed view of the recognition interface between H2BK120ub and BMI1. **f**, *In vitro* ubiquitination assays of wild-type and mutant cPRC1 complexes with unmodified or H2BK120ub-modified nucleosomes, analyzed by Coomassie blue staining (top) and Western blot (bottom). The middle row represents the ratio of the band intensity of H2A-ub (including H2A-ub and H2A-ub2) to that of H3, normalized to lane 2 (BMI1^WT^-NCP^H2BK120ub^, set to 1.00). The experiment was repeated at least three times with similar results.

To elucidate the structural mechanism by which H2BK120ub modification enhances PRC1 activity, we assembled human cPRC1 E3:E2∼ub enzyme complex bound to an H2BK120ub-modified nucleosome and determined its cryo-EM structure at an overall resolution of 2.8 Å (Fig. 1d, Extended Data Fig. 1a-g, and Supplementary Table 1). We observed that the cPRC1 E3:E2∼ub complex binds to the nucleosome surface in a manner similar to previously reported PRC1-nucleosome structures^37^, with the ubiquitin at Lys120 of histone H2B directly interacting with the RING domain of the PRC1 subunit BMI1 (Fig. 1d). The I44 patch (residues Leu8, Ile44, His68, and Val70) of H2BK120ub is in close proximity to a panel of hydrophobic residues of BMI1, and this interface is further stabilized by hydrogen-bonding interactions between the side chains of H2BK120ub^Arg42, Arg72^ and BMI1^Asp57^ (Fig. 1e and Extended Data Fig. 1h). The BMI1 residues involved in H2BK120ub recognition are highly conserved across species and in its close paralog MEL18/PCGF2. In contrast, other PCGF paralogs show variations in several of these residues, which might affect their association with H2BK120ub (Extended Data Fig. 2a). Substitution of BMI1^Asp57^ with the corresponding residues from other PCGFs led to a loss of the enhanced enzymatic activity of RNF2-BMI1 on the H2BK120ub nucleosome, suggesting that hydrogen-bonding interactions between H2BK120ub^Arg42,Arg72^ and BMI1^Asp57^ are essential for the histone crosstalk between H2BK120ub and cPRC1-mediated H2AK119ub deposition (Fig. 1f).

### Differential regulation of ncPRC1 variant activities by H2BK120ub

We next analyzed the impact of H2BK120ub on the catalytic activity of noncanonical PRC1 (ncPRC1) complexes. As shown in Fig. 2b, the RNF2-PCGF1 complex displayed stronger H2AK119 ubiquitination activity compared to the RNF2-PCGF6 complex, and this activity was remarkably enhanced by the incorporation of the ncPRC1-specific subunit RYBP, consistent with previous reports^11^. Interestingly, H2BK120ub stimulated the H2AK119 ubiquitination catalyzed by the RNF2-PCGF1 and RNF2-PCGF1-RYBP (referred to as ncPRC1.1) complexes, but had no effect on the activity of the RNF2-PCGF6 and RNF2-PCGF6-RYBP (referred to as ncPRC1.6) complexes (Fig. 2b). Cryo-EM analysis of the ncPRC1.1 complex associated with E2∼ub and the H2BK120ub nucleosome revealed that ncPRC1.1 assembles an E3:E2∼ub complex on the surface of nucleosome similar to that observed for the cPRC1 complex, with H2BK120ub contacting the RING domain of PCGF1, the counterpart of BMI1 (Fig. 2a, c, Extended Data Fig. 2b-g, and Supplementary Table 1). Structural superposition of the cPRC1 and ncPRC1.1 complexes revealed certain discrepancies between the interfaces of H2BK120ub with BMI1 and PCGF1, characterized by a noticeable shift in the position of H2BK120ub (Fig. 2d). Consequently, several contacting residues between H2BK120ub and PCGF1 differ from those observed between H2BK120ub and BMI1, which likely accounts for the variations in the residues involved in H2BK120ub association among PCGF paralogs (Fig. 2e and Extended Data Figs. 2a,h). In addition, a short helical turn followed by a loop structure in BMI1 (residues 110-124), which is absent in PCGF1, mediates direct interactions with the E2-conjugated ubiquitin, underscoring a notable structural difference between the catalytic modules of cPRC1 and ncPRC1.1 (Fig. 2d). Deletion of residues 110-124 in BMI1 moderately decreased the *in vitro* ubiquitination activity of the cPRC1 complex, suggesting that this region-mediated interaction with the E2-conjugated ubiquitin contributes to the E3 ligase activity of cPRC1 (Fig. 1f).

**Figure 2.**
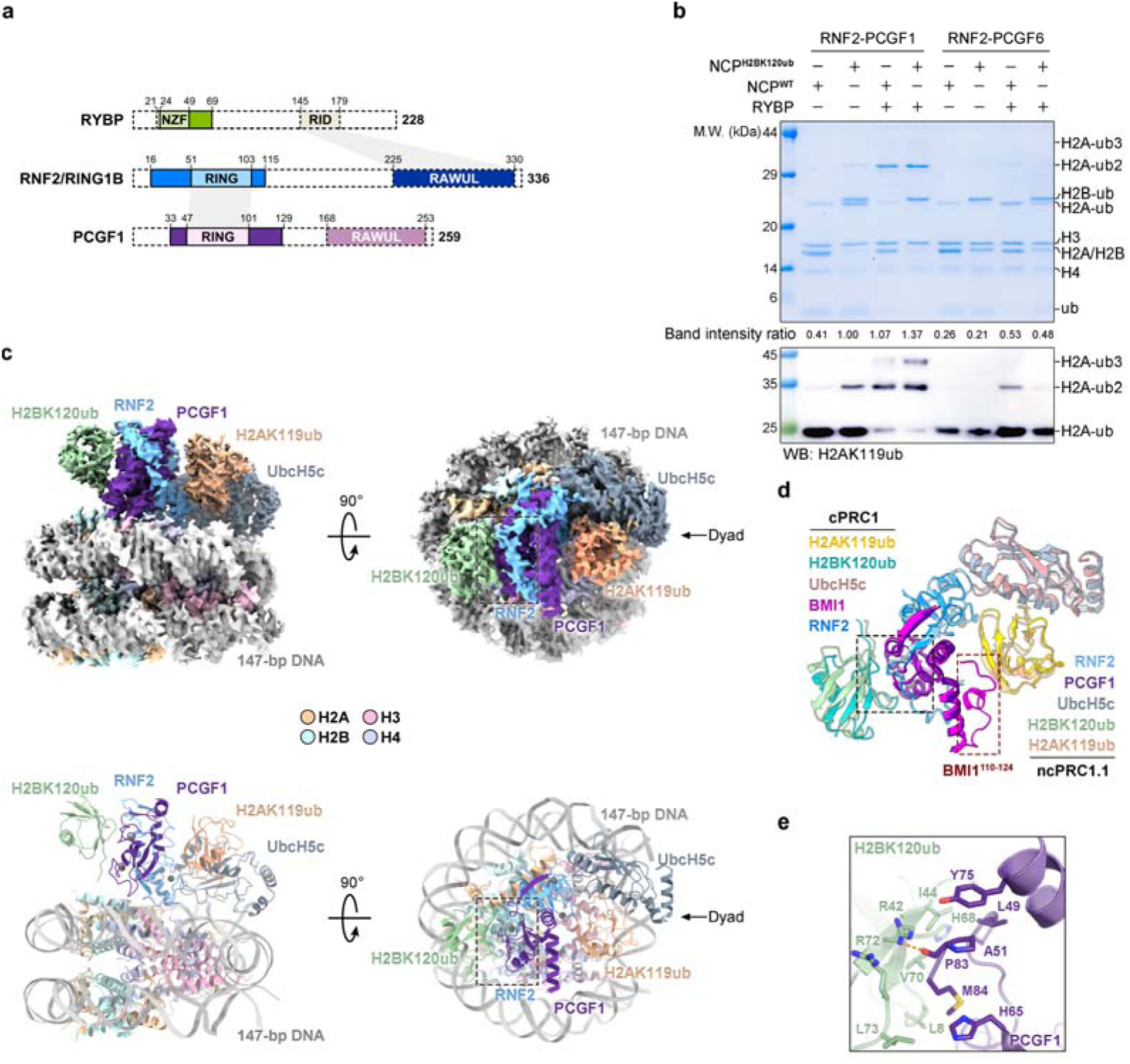
H2BK120ub preferentially interacts with ncPRC1.1 complex, rather than ncPRC1.6, to promoting the deposition of H2AK119ub on nucleosomes. **a**, Schematic of the domain organizations of the human ncPRC1.1 ternary complex. Abbreviations of domain names: NZF, Npl4-type zinc finger; RID, Ring1B-interacting domain; RAWUL, ring finger and WD40 associated ubiquitin-like. **b**, *In vitro* ubiquitination assays of ncPRC1.1 and ncPRC1.6 with unmodified or H2BK120ub-modified nucleosomes, analyzed by Coomassie blue staining (top) and Western blot (bottom). The middle row represents the ratio of the band intensity of H2A-ub (including H2A-ub, H2A-ub2, and H2A-ub3) to that of H3, normalized to lane 2 (RNF2-PCGF1-NCP^H2BK120ub^, set to 1.00). The experiment was repeated at least three times with similar results. **c**, Overall structure of human ncPRC1.1-E2∼ub-NCP^H2BK120ub^ complex. Cryo-EM density map (upper panel) and atomic model (lower panel) of ncPRC1.1-E2∼ub-NCP^H2BK120ub^ complex are shown from two orthogonal views. The cryo-EM map is segmented according to the components of ncPRC1.1-E2∼ub-NCP^H2BK120ub^ complex. The color scheme of ncPRC1.1-E2∼ub-NCP^H2BK120ub^ complex is the same as depicted in (a), and the 147-bp DNA chains are shown in light and dark gray, respectively. Ub, ubiquitin; E2, UbcH5c. **d**, Structural comparison of cPRC1-E2∼ub-NCP^H2BK120ub^ and ncPRC1.1-E2∼ub-NCP^H2BK120ub^. **e**, Detailed view of the recognition interface between H2BK120ub and PCGF1.

### H2BK120ub modulates the competitive binding of ncPRC1.4 subunits to the acidic patch of nucleosomes

Specific recognition of H2AK119ub-modified nucleosomes by the ncPRC1 subunit RYBP is crucial for recruiting ncPRC1 complexes to chromatin and facilitating the propagation of H2AK119ub marks to neighboring nucleosomes through a positive-feedback loop^10,12^. Previous studies have shown that the NZF domain of RYBP (RYBP^NZF^) interacts simultaneously with the ubiquitin moiety on H2AK119 and the acidic patch of nucleosomes, outcompeting the binding of the ncPRC1.4 (RNF2-BMI1-RYBP) catalytic module to nucleosomes^16,17^. As H2BK120ub can directly bind to the catalytic module of both cPRC1 and ncPRC1.1 complexes to enhance their catalytic activities as demonstrated above, we next investigated whether the presence of H2BK120ub on nucleosomes influences the manner in which ncPRC1 complexes associate with their H2AK119ub nucleosome targets.

To examine whether H2BK120ub alters the engagement of human ncPRC1.4 complex (containing the BMI1 subunit, as in cPRC1) with H2AK119ub-modified nucleosomes, we determined cryo-EM structures of ncPRC1.4 bound to nucleosomes bearing both H2BK120ub and H2AK119ub modifications (NCP^H2BK120ub&H2AK119ub^) or H2AK119ub alone (NCP^H2AK119ub^), revealing distinct binding modes (Fig. 3a). Masked 3D classification of the EM densities for ncPRC1.4 showed that in 63% of the dataset, two ncPRC1.4 complexes are anchored to opposite sides of a nucleosome, with the catalytic module (RNF2-BMI1) binding to the nucleosomal acidic patch. H2BK120ub associates with BMI1 in a manner similar to that observed in the cPRC1-E2∼ub-NCP^H2BK120ub^ complex structure. In the remaining portion of the dataset, a single ncPRC1.4 complex binds to one side of a nucleosome through its catalytic module interacting with H2BK120ub and the acidic patch, while its RYBP^NZF^ domain engages with the acidic patch on the opposite side of the nucleosome (Fig. 3a, Extended Data Fig. 3, and Supplementary Table 1). In comparison, when associated with nucleosomes containing only H2AK119ub modifications, ncPRC1.4 complexes exhibited more heterogeneous binding patterns: in 68% of the cryo-EM dataset, the RYBP^NZF^ domain occupied the nucleosome surface; in 19% of the dataset, the catalytic module and the RYBP^NZF^ domain were positioned on opposite sides of the nucleosome; and in 13%, only the catalytic module was bound to one side of the nucleosome (Fig. 3a, Extended Data Fig. 4, and Supplementary Table 1).

**Figure 3.**
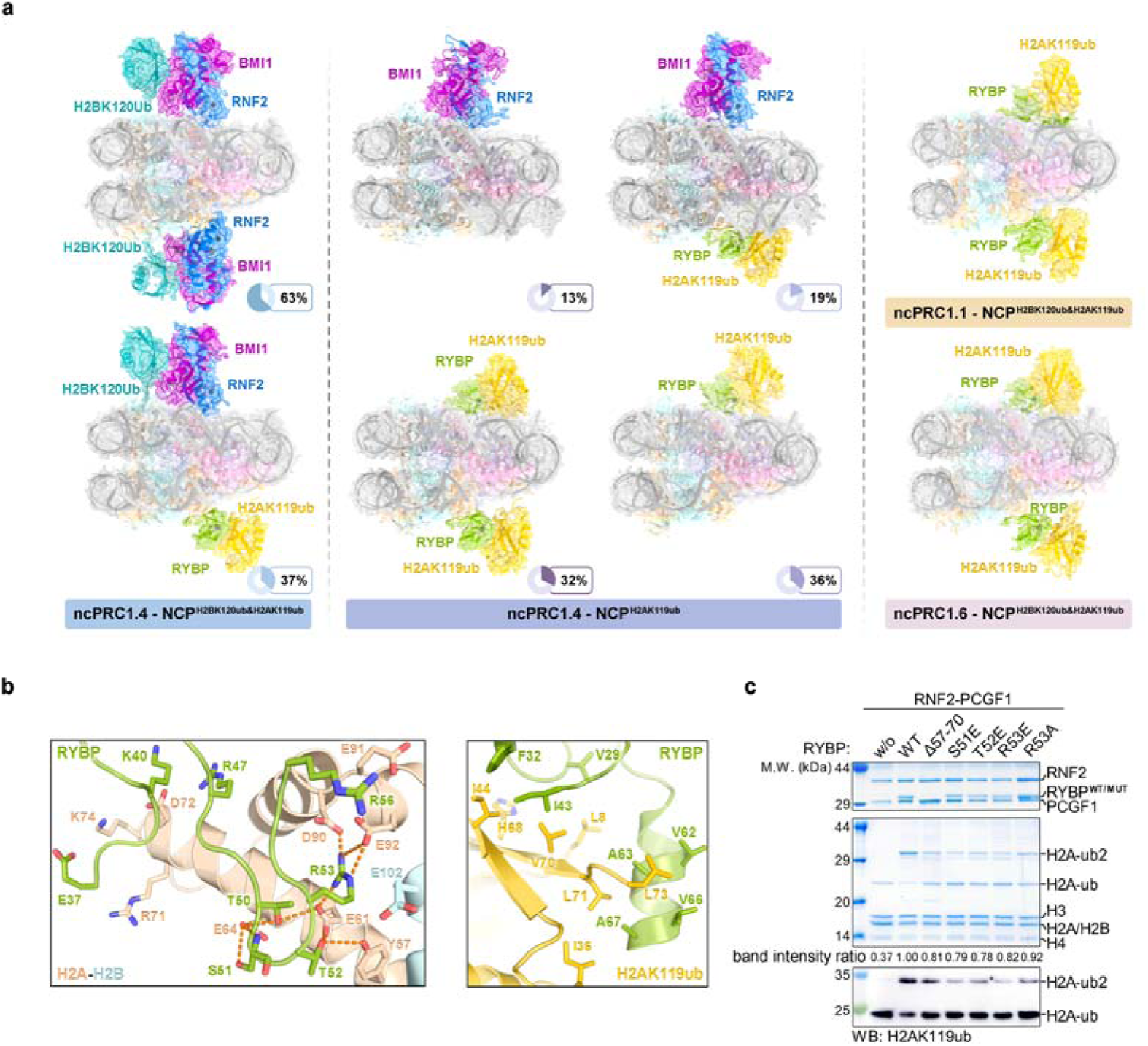
H2BK120ub alters the binding pattern of the ncPRC1.4 complex to H2AK119ub-modified nucleosomes. **a**, Cryo-EM density maps and atomic models of human ncPRC1.4-NCP^H2BK120ub&H2AK119ub^ complex (left), human ncPRC1.4-NCP^H2AK119ub^ complex (middle), human ncPRC1.1-NCP^H2BK120ub&H2AK119ub^ complex (top right), and human ncPRC1.6-NCP^H2BK120ub&H2AK119ub^ complex (bottom right) in the dyad view of nucleosome. **b**, Detailed view of the recognition interfaces between RYBP and the nucleosome acidic patch (left), and between RYBP and H2AK119ub (right). **c**, *In vitro* ubiquitination assays of wild-type and mutant ncPRC1.1 complexes with unmodified nucleosomes, analyzed by Coomassie blue staining (middle) and Western blot (bottom). The upper panel shows the protein purity of wild-type and mutant ncPRC1.1 complexes loaded at a higher concentration than in the ubiquitination assay. The middle row represents the ratio of the band intensity of H2A-ub (including H2A-ub and H2A-ub2) to that of H3, normalized to lane 2 (RNF2-PCGF1^WT^-NCP^H2BK120ub^, set to 1.00). The experiment was repeated at least three times with similar results. w/o, without RYBP.

These observations indicate that the presence of H2BK120ub remarkedly alters the binding patterns of ncPRC1.4 on H2AK119ub-modified nucleosomes. The direct interaction between H2BK120ub and BMI1 enhances the binding affinity of the catalytic module to nucleosome, enabling it to outcompete the RYBP^NZF^ domain for nucleosome association.

### ncPRC1.1 and ncPRC1.6 associate with H2AK119ub-modified nucleosomes in an H2BK120ub-insensitive manner

We further investigated whether H2BK120ub regulates the competitive binding between the catalytic module and the RYBP^NZF^ domain to nucleosomes in ncPRC1.1 and ncPRC1.6 complexes. Cryo-EM structures of ncPRC1.1 or ncPRC1.6 associated with NCP^H2BK120ub&H2AK119ub^ demonstrated that the RYBP^NZF^ domain is sandwiched between H2AK119ub and the acidic patch of nucleosome, while the EM density for H2BK120ub is barely visible (Fig. 3a, Extended Data Fig. 5a-f, Extended Data Fig. 6, and Supplementary Table 1). In these structures, the interfaces of RYBP^NZF^ with the I44 patch of H2AK119ub and the acidic patch of nucleosome closely resemble those observed in the previously reported ncPRC1.4-NCP^H2AK119ub^ structures^16,17^ (Fig. 3b and Extended Data Fig. 5g). Additionally, an α-helix immediately C-terminal to RYBP^NZF^ makes contact with the I36 patch (residues Ile36, Leu71, and Leu73) of H2AK119ub (Fig. 3b and Extended Data Fig. 5g). Mutations in RYBP (S51E, T52E, R53E, and R53A), which disrupt the interaction with the acidic patch of the nucleosome, noticeably impaired the *in vitro* H2AK119ub activity of the ncPRC1.1 complex. Deletion of the interacting RYBP α-helix (Δ57-70) also led to a modest reduction in this activity (Fig. 3c).

Although H2BK120ub can bind to PCGF1 in the ncPRC1.1-E2∼ub-NCP^H2BK120ub^ complex structure, this interaction does not confer a similar competitive advantage for the catalytic module of ncPRC1.1 when binding to NCP^H2BK120ub&H2AK119ub^. This discrepancy between ncPRC1.1 and ncPRC1.4 complexes may arise from differences in the binding affinities of H2BK120ub to various PCGF paralogs, as well as variations in the extent to which these PCGFs interact with the nucleosomal acidic patch.

### H2BK120ub and H2AK119ub colocalize in gene bodies of a group of developmental genes in mESCs

To analyze the effects of the H2BK120ub modification on the genome-wide distribution of H2AK119ub in mouse embryonic stem cells (mESCs), we generated an *Rnf20* knockout cell line using the CRISPR/Cas9 system (Extended Data Fig. 7a). Unexpectedly, the loss of *Rnf20* expression not only substantially abolished the H2BK120ub marks it catalyzes, but also obviously reduced the total levels of Bmi1 and H2AK119ub (Extended Data Fig. 7a). We treated the mESCs with the ribosome inhibitor cycloheximide (CHX) to assess Bmi1 protein stability, which showed that Bmi1 protein levels remained stable over time in wild-type mESCs, whereas in *Rnf20* knockout cells, Bmi1 protein levels progressively declined (Extended Data Fig. 7b). Additionally, L-azidohomoalanine (AHA) pulse-labeling of newly synthesized proteins revealed that the levels of Bmi1 incorporated with AHA were comparable between wild-type and *Rnf20* knockout cells (Extended Data Fig. 7c). These results suggest that Rnf20 knockout promotes the degradation of Bmi1 without affecting its synthesis. To compensate for the reduced Bmi1 protein levels, we transiently expressed Flag-tagged Bmi1 in *Rnf20* knockout cells, restoring Bmi1 levels similar to those in wild-type mESCs for subsequent analysis (Extended Data Figs. 7d,e).

We first assessed the enrichment levels of H2AK119ub and H2BK120ub in these mESCs (Fig. 4a and Extended Data Fig. 7f,g). Consistent with previous studies, we observed that H2AK119ub was predominantly enriched at promoter regions, whereas H2BK120ub was more concentrated in gene bodies. Notably, we identified a subset of genes exhibiting high enrichment of both H2AK119ub and H2BK120ub in gene bodies. Following *Rnf20* knockout and Bmi1 restoration, the gene body enrichment of both H2AK119ub and H2BK120ub at these genes was abolished, while the enrichment of Bmi1 remained unaltered (Fig. 4a and Extended Data Fig. 7h). This suggests that the depletion of H2AK119ub was not due to the loss of PRC1 distribution, but rather to decreased enzymatic activity of PRC1. This is consistent with our previous data showing that H2BK120ub enhances PRC1 activity, facilitating the deposition of H2AK119ub.

**Figure 4.**
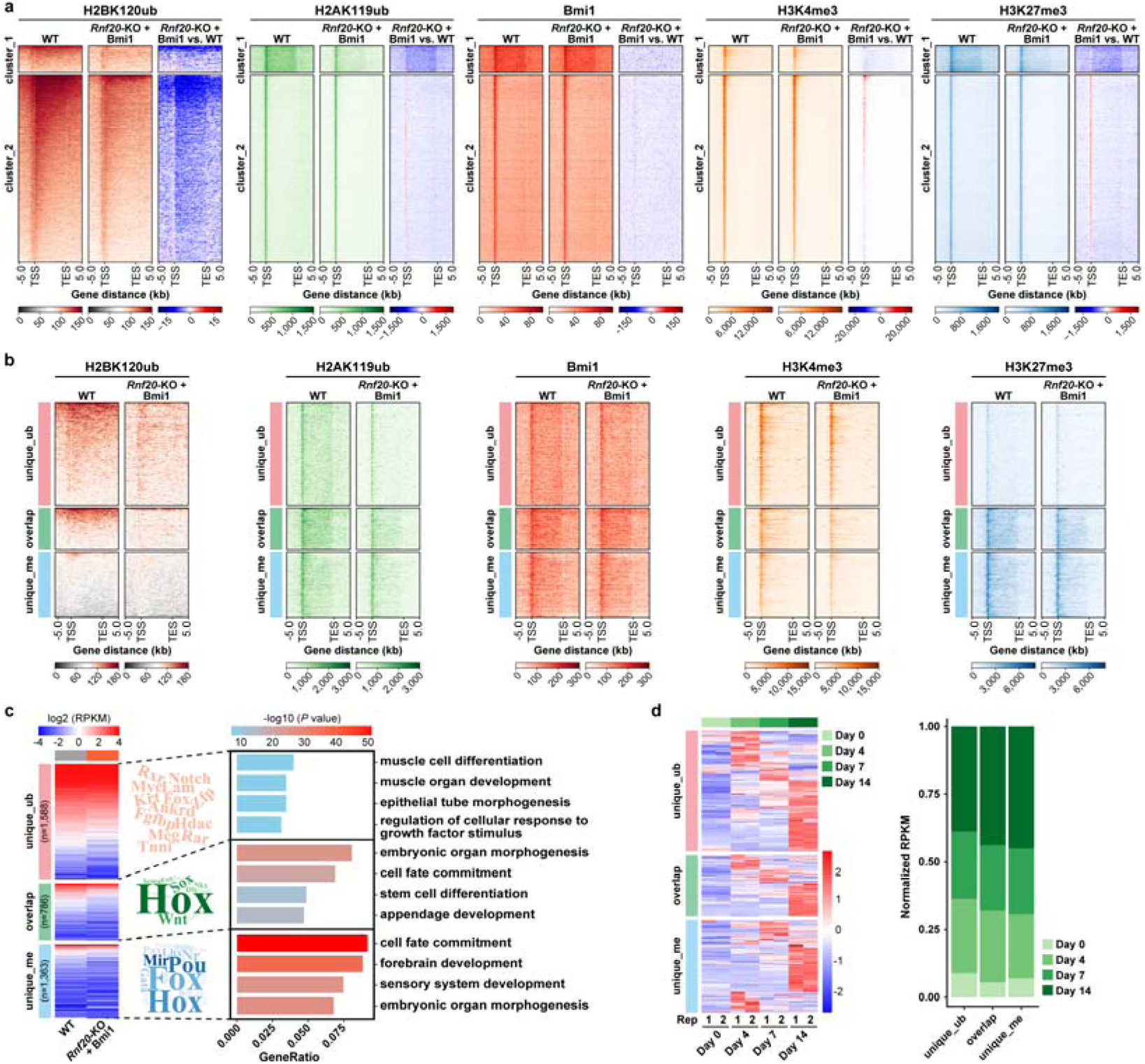
H2BK120ub and H2AK119ub colocalize in a subset of developmental genes in mESCs. **a**, Heatmaps show normalized H2BK120ub, H2AK119ub, Bmi1, H3K4me3, and H3K27me3 signals in WT and *Rnf20*KO+Flag-Bmi1 mESCs. Signal differences were calculated by subtracting WT mESC signals from those of *Rnf20*-KO+ Flag-Bmi1. Data were plotted across 5 kb windows spanning the TSS to TES of all genes determined by UCSC Genes 2013. Genes were clustered into two groups using K-Means based on WT H2BK120ub and H2AK119ub signals. Heatmaps were sorted by WT H2BK120ub enrichment. TSS, transcription start site. TES, transcription end site. **b**, Heatmaps of normalized H2BK120ub, H2AK119ub, Bmi1, H3K4me3, and H3K27me3 signals in different gene groups from WT and *Rnf20-*KO+Flag-Bmi1 mESCs, shown similarly as in Fig. 4a. **c**, Heatmaps (left) show gene expression in different gene groups from WT and *Rnf20-*KO+Flag-Bmi1 mESCs. Word clouds (medium) illustrate the gene families in each group. GO analysis (right) presents the top four GO terms ranked by *P* values, with *P* values calculated using the hypergeometric test. **d**, Heatmaps (left) display normalized gene expression and a percentage stacked bar plot (right) shows the distribution of average normalized expression in unique_ub, overlap, and unique_me from WT mESCs at differentiation time points of 0, 4, 7, and 14 days. The normalization was applied by rows.

Compared to the well-characterized bivalent histone modifications H3K4me3 and H3K27me3, the co-occurrence of H2AK119ub and H2BK120ub at genes represents a different type of bivalency that integrates both repressive and active histone marks. Although some overlap was observed, the genes marked by both H2AK119ub and H2BK120ub were largely distinct from those associated with H3K4me3 and H3K27me3 (Fig. 4b,c and Extended Data Fig. 7i). We designated the former as ubiquitination-bivalent genes and the latter as methylation-bivalent genes for further comparison. The unique ubiquitination-bivalent genes were marked with high levels of H3K4me3 and relatively low enrichment of H3K27me3. Upon *Rnf20* knockout and Bmi1 restoration, the H3K27me3 levels at these gene bodies were further reduced, likely due to the decrease in H2AK119ub levels (Fig. 4b). Ubiquitination-bivalent genes were linked to muscle development, epithelial tube morphogenesis, and cellular response to growth factor stimuli, while methylation-bivalent genes corresponded to more general developmental regulation (Fig. 4c). Notably, the expression levels of unique ubiquitination-bivalent genes were higher than those of unique methylation-bivalent genes. These genes also exhibited considerably higher rates of RNA synthesis and higher levels of RNA Pol II at their promoters compared to their gene bodies (Extended Data Figs. 7j,k). These observations suggest that ubiquitination-bivalent genes are primed for activation and may be activated more rapidly than methylation-bivalent genes. Further analysis of gene expression patterns from available mESC differentiation datasets^38^ revealed that ubiquitination-bivalent genes were indeed activated earlier than methylation-bivalent genes (Fig. 4d). Together, these data suggest that H2BK120ub promotes the deposition of H2AK119ub at gene bodies, forming ubiquitination-bivalent genes that can be quickly activated during mESC differentiation.

## Discussion

The intrinsic interplay between PcG and TrxG complexes in orchestrating chromatin states has long been recognized as a fundamental mechanism of epigenetic regulation. Beyond the traditional antagonism between these two complexes, we present here an unexpected histone crosstalk involving the stimulatory effects of the active histone mark H2BK120ub on PRC1 activities, suggesting a novel role for PRC1 in bridging the functions of PcG and TrxG complexes. Characterization of the ubiquitination-bivalent genes in mESCs, distinct from the classical methylation bivalency, further highlights the potential roles of this histone crosstalk in development regulation.

In this study, we provide structural elucidation of the direct association between H2BK120ub and cPRC1 as well as ncPRC1.1 on the surface of nucleosomes, demonstrating how this interaction enhances PRC1 activities (Fig. 5). H2BK120ub, primarily associated with transcription elongation, is enriched in the gene bodies of actively transcribed genes and facilitates the efficient methylation of histone H3K4 and H3K79 for gene activation^21,22,39^. Notably, H2BK120ub employs a mechanism analogous to its regulation of PRC1 by directly interacting with the H3K4 methyltransferase MLL/COMPASS and the H3K79 methyltransferase DOT1L to boost their enzymatic activities^23,40^. The acidic patch, a nucleosome surface commonly targeted by chromatin modulators, mediates the binding of PRC1, COMPASS, and DOT1L to nucleosomes, with H2BK120ub further strengthening these associations in a synergistic manner^23,40^. In contrast, H2BK120ub alone can partially block the acidic patch, thereby interfering with the binding of some other chromatin modulators, such as RCC1, Sir3, and LANA, to this site^41^. This context-dependent dual regulation of nucleosome accessibility by H2BK120ub may broadly impact the interactions between various chromatin modulators and nucleosomes, requiring further exploration of these complex dynamics.

**Figure 5.**
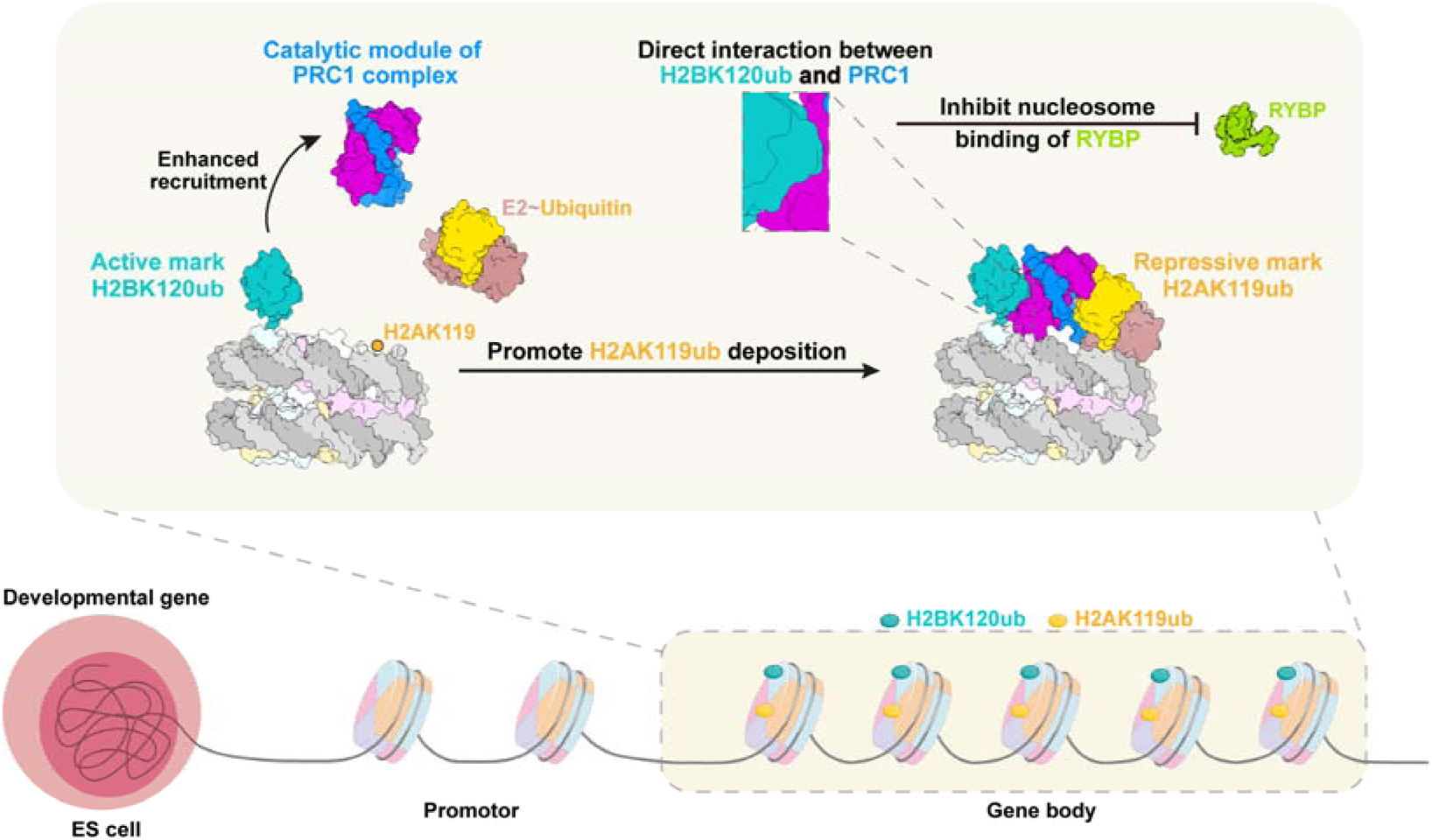
A working model of PRC1 complex action on nucleosomes in the presence of H2BK120ub. Direct interaction between H2BK120ub and specific PCGF subunits promotes PRC1 activity while limiting RYBP association with nucleosomes. Co-enrichment of H2BK120ub and H2AK119ub is observed in the gene bodies of a subset of developmental genes.

In addition to modulating the enzymatic activities of cPRC1 and ncPRC1.1, H2BK120ub also affects how the catalytic module and the RYBP subunit of the ncPRC1.4 complex compete for binding to the acidic patch of nucleosomes (Fig. 5). The concurrent binding of RYBP and its close homolog YAF2 to both the ubiquitin and the acidic patch of H2AK119ub-modified nucleosomes promotes the propagation of H2AK119ub depositions to neighboring nucleosomes through a positive-feedback mechanism^10–12^. Previous cryo-EM studies have revealed that RYBP can outcompete the catalytic module for access to the acidic patch when H2AK119ub is present on the nucleosome^16,17^. Our analysis of ncPRC1.1, a key complex for H2AK119ub spreading, revealed that the addition of H2BK120ub alongside H2AK119ub did not alter RYBP’s binding preference for the nucleosome surface, suggesting that H2BK120ub does not impact ncPRC1.1 function in H2AK119ub propagation. Similar observations were made for ncPRC1.6. However, in the case of the ncPRC1.4 complex, H2BK120ub notably altered the binding pattern, favoring the association of the catalytic module BMI1-RNF2 with the nucleosome acidic patch over that of RYBP. Despite sharing the well-characterized BMI1 subunit with cPRC1, the specific functions of ncPRC1.4 remain poorly understood. The altered nucleosome-binding mode adopted by ncPRC1.4 confines the catalytic module to a single nucleosome, thereby attenuating RYBP-dependent recognition of H2AK119ub and restricting the propagation of this repressive mark to adjacent nucleosomes. Moreover, even in the absence of H2BK120ub, BMI1-RNF2 partially occupies the acidic patch in certain 3D classes of the cryo-EM dataset, suggesting that it may have a higher nucleosome-binding affinity than other PCGF paralogs. Given that BMI1-RNF2 does not exhibit stronger enzymatic activity than other PCGF-RNF2 paralogs^11^, its preference for acidic patch occupation may contribute to PRC1 regulation through non-enzymatic mechanisms.

We found that the histone crosstalk between H2BK120ub and H2AK119ub is functionally associated with a subset of development-related genes that are primed for earlier activation compared to the canonical methylation-bivalent genes in mESCs (Fig. 5). PRC1 has been shown to modulate active transcription in a multifaceted, context-dependent manner rather than solely acting as a transcriptional repressor. Studies in both *Drosophila* and mammals indicate that PRC1 can be recruited to specific gene subsets lacking H3K27me3 to mediate transcription activation involved in key developmental processes and tumorigenesis^42–47^. In this work, we establish a direct connection between PRC1 and active transcription through its association with the activating mark H2BK120ub. The gene body localization of H2BK120ub drives a similar distribution pattern for both Bmi1 and H2AK119ub that diverges from their typical promoter enrichment, although the specific functions of PRC1 and H2AK119ub within gene bodies remain to be elucidated. Moreover, these ubiquitination-bivalent genes, which largely differ from canonical methylation-bivalent genes, typically exhibit low H3K27me3 levels and are more readily activated during embryogenesis. This observation aligns with previous findings that the transcription activation function of PRC1 is independent of PRC2^42–44^.

In summary, our structural elucidation of H2BK120ub-mediated stimulation of PRC1 activity provides mechanistic insights into how PRC1 presumably orchestrates the balance between gene silencing and activation. These findings are essential for understanding the regulation of PRC1 on poised chromatin states and lineage specification in embryonic stem cells. Our work also establishes a structural framework for further investigating PRC1 dysregulation, particularly BMI1 overexpression in cancer stem cell maintenance and tumorigenesis, and contributes to the development of epigenetic therapies for cancer.

## Acknowledgements

We are grateful to the staff of the core facilities at Shanghai Institute of Precision Medicine and the National Facility for Protein Science in Shanghai (NFPS), Zhangjiang Lab for their instrument support and technical assistance. We also thank our college Dr. Shiyan Yu for his insightful suggestions on our studies. This work was supported by the National Key R&D Program of China (2022YFA1302800), the National Natural Science Foundation of China (32022036), and the Innovative Research Team of High-level Local Universities in Shanghai (SHSMU-ZLCX20212300).

## Author contributions

Y.Y., D.C., and G.B. performed sample preparation, EM data collection, and biochemical analyses; Y. Z. performed EM data analyses and helped with manuscript preparation; X.W. and Y.L. conducted analyses related to mESCs; F.W. and J.C. helped with bioinformatic analyses; Z.T., J.N., and H.X. helped with biochemical analyses; M.L. provided valuable advice on the project and the manuscript; J.H. designed and supervised the research; J.H., D.F., C.Y. and Z.Z. wrote the manuscript with help from all other authors.

## Author information

The authors declare no competing financial interests. Correspondence and requests for materials should be addressed to J.H.

## METHODS

### Protein Expression and Purification

Human canonical PRC1.4 (cPRC1.4) core complex, which includes RNF2 and BMI1, was cloned into a modified pMLink vector with an N-terminal 6×His-3×Flag tag followed by a 3C protease cleavage site. Human noncanonical PRC1.1 core complex (RNF2, PCGF1, and RYBP), noncanonical PRC1.4 core complex (RNF2, BMI1, and RYBP), and noncanonical PRC1.6 core complex (RNF2, PCGF6, and RYBP) were cloned into the same vector.

For the preparation of the cPRC1.4 core complex, RNF2 and BMI1 were co-expressed in Expi293 cells (Gibco) and were purified with Anti-DYKDDDDK G1 Affinity Resin (GenScript) in Lysis Buffer A (50 mM Tris-HCl pH 7.5, 150 mM NaCl, 10% glycerol, 1 mM DTT, 1 mM PMSF, 5 mM benzamidine, 1 µg/ml leupeptin, and 1 µg/ml pepstatin). The 6×His-3×Flag tag was removed through 3C protease digestion, and the complex was further purified through Superdex 200 Increase gel-filtration chromatography (GE Healthcare) in Column Buffer A (25 mM HEPES pH 7.5 and 150 mM NaCl). The ncPRC1.1, ncPRC1.4, and ncPRC1.6 core complexes were prepared following the same procedure.

UbcH5c∼ub was prepared by conjugating the purified UbcH5c bearing the C85S and S22R mutations with ubiquitin *in vitro*. Briefly, 6×His-tagged UbcH5c and 6×His-SUMO-tagged ubiquitin were expressed in *E. coli* Transetta (DE3) and purified with Ni-NTA agarose beads (Qiagen) in Lysis Buffer B (50 mM Tris-HCl pH 7.5, 500 mM NaCl, 10% glycerol, 5 mM 2-mercaptoethanol, 1 mM PMSF, 5 mM benzamidine, 1 μg/ml leupeptin, and 1 μg/ml pepstatin), respectively. The affinity tags were removed through 3C protease or Ulp1 protease digestion, followed by HiLoad 75 Increase gel-filtration chromatography (GE Healthcare) in Column Buffer B (25 mM Tris-HCl pH 7.5 and 150 mM NaCl). The purified ubiquitin was further processed by rebinding to Ni-NTA agarose beads, followed by Superdex 75 Increase gel-filtration chromatography (GE Healthcare) in Column Buffer B. Then, UBA1, UbcH5c and ubiquitin were mixed at a molar ratio of 1:15:45 in Column Buffer B supplemented with 2 mM ATP, and incubated at 30 °C for 24 h. The UbcH5c∼ub conjugate was further purified through Superdex 75 Increase gel-filtration chromatography (GE Healthcare).

### Nucleosome Preparation

Nucleosomes containing H2BK120ub were prepared as described previously^48,49^. In brief, a histone H2B_K120C and a ubiquitin_G76C were introduced to mediate the chemical cross-linking between residue K120 of histone H2B and the C terminus of ubiquitin. *X. laevis* histones, including H2A, H2B, H3, H4 and H2B_K120C were expressed in *E. coli* BL21 (DE3) and isolated as inclusion bodies. After sequential purifications through Q Sepharose HP (GE Healthcare) and SP Sepharose HP (GE Healthcare) columns in denaturing buffer (20 mM Tris-HCl pH 7.5, 8 M urea, 0-0.5 M NaCl, 1 mM EDTA, and 5 mM 2-mercaptoethanol), the histone proteins were dialyzed thoroughly against distilled water containing 2 mM 2-mercaptoethanol, and were then lyophilized and stored at -80 °C.

The Ub_G76C was fused with a 6×His tag at the N-terminus following a 3C protease cleavage site (6×His-3C-Ub_G76C) and was purified with Ni-NTA agarose beads and HiLoad Superdex 75 gel-filtration chromatography. After thorough dialysis against 50 mM ammonium acetate (pH 4.5), the ubiquitin proteins were lyophilized and stored at -80 °C. The histone H2B_K120C and the 6×His3C-Ub_G76C proteins were cross-linked by the chemical reagent dichloroacetone (DCA). The H2B-Ub conjugate was sequentially purified through a denatured Ni-NTA affinity purification and semi-preparative C4 reversed-phase high-performance liquid chromatography, and the 6×His-3C tag was removed through 3C protease digestion. The purified H2B-Ub proteins were lyophilized and stored at -80 °C.

The NCP^H2BK120ub^ was reconstituted from four core histones (H3, H4, H2A and H2B-Ub) and the 147-bp Widom 601 DNA. First, histone octamers were assembled and separated through HiLoad Superdex 200 gel filtration in refolding buffer (20 mM Tris-HCl pH 7.5, 2 M NaCl, 1 mM EDTA, and 5 mM 2-mercaptoethanol). Then, the purified histone octamer was mixed with the Widom 601 DNA at a molar ratio of 0.9:1 and was dialyzed against reconstitution buffer (10 mM Tris-HCl pH 7.5, 1 mM EDTA, 1 mM DTT, and 0.25-2 M KCl). The reconstituted NCP_H2BK120ub was further purified through HiLoad Superdex 200 gel filtration, and stored in small aliquots at -80 °C. The unmodified NCP was *in vitro*-reconstituted similarly as described for the NCP^H2BK120ub^ reconstitution. NCP^H2BK120ub&H2AK119ub^ and NCP^H2AK119ub^ were obtained through *in vitro* ubiquitination reactions as described below, followed by purification using Superose 6 Increase gel-filtration chromatography.

### Cryo-EM sample preparation and data collection

The cPRC1.4 core complex and the ncPRC1.1 core complex were each mixed with the H2BK120ub nucleosome and the UbcH5c∼ub conjugate at a molar ratio of 16:1:64, and incubated at 4 °C for 1 h. The ncPRC1.1, ncPRC1.4 and ncPRC1.6 core complexes were each mixed with NCP^H2BK120ub&H2AK119ub^ or NCP^H2AK119ub^ at a molar ratio of 16:1, and incubated at 4 °C for 1 h. These mixtures were then separately fractionated using the GraFix method^50^. The samples were loaded onto a continuous 10-30% gradient of glycerol with a 0-0.1% gradient of glutaraldehyde in GraFix Buffer (25 mM HEPES pH7.5 and 150 mM NaCl) generated with a BioComp gradient master, and were subjected to ultracentrifugation at 37,000 rpm at 4 °C for 15 h, using a SW60 Ti rotor (Beckman). The fractions containing the desired complex were added with 50 mM Tris-HCl pH 7.5, dialyzed against GraFix Buffer, and concentrated to approximately 200 ng/μl as determined by UV absorbance at 260 nm. 3 μl of the samples were applied onto glow-discharged holey carbon grids (Quantifoil R1.2/1.3, Au 300 mesh, Cu 200 mesh). The grids were blotted for 2.5 s, and were then plunged into liquid ethane cooled by liquid nitrogen, using a Vitrobot Mark IV (FEI).

The samples were observed under a Titan Krios G3i transmission electron microscope (FEI) operated at 300 kV. The images were collected on a K3 direct electron detector (Gatan) with a pixel size of 1.10 Å and a defocus range of −1.2 to −1.6 μm. Each micrograph was dose-fractioned to 32 frames with 0.1 s exposure time for each frame. The total accumulated dose of each micrograph was 50.0 e^-^/Å^2^. The imaging conditions are also listed in Supplementary Table 1.

### Cryo-EM data analysis and model building

Motion correction was applied to the dose-fractioned cryo-EM image stacks using MotionCor2 with dose-weighting^51,52^. The contrast-transfer function (CTF) parameters of each image were determined with CTFFIND4.1^53^, and automatic particle-picking was carried out using Topaz^54^. 2D classification, 3D classification, 3D auto-refinement, CTF refinement and Bayesian polishing were performed with RELION-4.0^55^. An overview of the data processing procedure is shown in Extended Data Figs. 1a, 2b, 3a, 4a, 5a, and 6a. After 1 round of 2D classification and 1-2 rounds of 3D classification with exhaustive angular searches, particles were processed with 3D auto-refinement and solvent-masked post-processing, which generated a cryo-EM density map with an overall resolution. To improve the local map density, the particles were further processed through masked 3D classifications with partial signal subtraction^56^. The resolution estimation was based on the gold-standard Fourier shell correlation (FSC) 0.143 criterion, and the local resolution was estimated with ResMap^57,58^.

Model building was carried out by fitting the available structures of PRC1-UbcH5c-NCP and H2BK120ub-NCP (PDB codes: 4R8P and 6KIU) in the EM density maps using UCSF Chimera^59^. The models were then manually built in Coot^60^, and real-space-refined with secondary structure restraints in Phenix^61^.

### *In vitro* ubiquitination assays

For a 20-μl *in vitro* ubiquitination reaction system, 0.48 μM wild-type or mutant cPRC1.4 core complexes, 0.03 μM UBA1, 0.48 μM UbcH5c, 10 μM Ub, 3 mM ATP, and 1 μM nucleosomes were mixed in a buffer containing 50 mM Tris-HCl pH7.5, 75 mM NaCl, 2 mM MgCl_2_, 10 μM ZnCl_2_, and 1 mM DTT. For noncanonical PRC1 complexes, 0.09 μM E3 and 0.09 μM UbcH5c were added. After incubation at 37 L for 1 h, 5 μl of 5×SDS loading buffer was added to stop the reaction. The samples were then loaded onto a 13.5% SDS-PAGE gel and analyzed by Coomassie staining and Western blot.

### Cell culture and cell lines

HEK Expi293 cells adapted for suspension were cultured in Union293 medium (Union-Biotech) at 37 °C, 5% CO_2_, and 110 rpm in a humidified shaker. The E14 mESC line was cultured on 0.1% gelatin-coated dishes in DMEM supplemented with 15% fetal bovine serum (FBS), 1% antibiotic solution (penicillin/streptomycin), 1% GlutaMAX, 1% MEM nonessential amino acids, 1% sodium pyruvate, 0.1 mM 2-mercaptoethanol, and 1000 U/ml recombinant leukemia inhibitory factor (LIF) at 37°C and 5% CO_2_ in a humidified incubator. *Rnf20*-KO cell lines were generated using sgRNAs cloned into the pSpCas9(BB)-2A-Puro plasmid from Dr. Feng Zhang’s lab (Addgene plasmid #48139).

### Protein degradation

Protein degradation was performed as previously described^62^. In brief, WT and *Rnf20*-KO mESCs were seeded into 6-well plates with an equal number of cells in each well. After 24 h, 25 μg/ml of cycloheximide (Aladdin, Cat. #C112766) was added into each well, and cells were collected in SDS loading buffer at 0, 2, 5, and 10 h after drug treatment.

### L-azidohomoalanine (AHA) labeling

WT and *Rnf20-*KO mESCs were seeded into 10 cm dishes with equal numbers of cells. After 24 h, the cells were switched to medium lacking methionine and cultured for 1 h. Then, they were cultured for an additional 2 h in methionine-free medium supplemented with 500 µM AHA (MedchemExpress, Cat. #HY-140346A). Cells were collected, washed twice with PBS, and lysed with 1 ml of RIPA buffer (50 mM Tris-HCl pH 7.5, 150 mM NaCl, 1% NP-40, 0.5% sodium deoxycholate, 0.1% SDS, 1 mM EDTA, and 1×protease inhibitor). The lysates were dounced with 50 strokes and rotated at 4 °C for 30 min. Samples were centrifuged at 12,000 g for 15 min at 4 °C. The supernatants (800 µl) were collected and mixed with the clicking buffer (8 µl of 100 mM Biotin-Alkyne, 24 µl of 5 mM Cu(II)-BTTAA, and 8 µl of 500 mM Ascorbic Acid), and incubated at 30 °C for 1 h. To quench the reaction, 0.8 µl of 100 mM Bovine Calf Serum was added to the samples, which were incubated at 30 °C for an additional 15 min. The protein was then purified using the methanol precipitation method. 5% of the purified protein was taken as input and boiled with SDS loading buffer. Pre-washed streptavidin beads (Smart-lifesciences, Cat. #SM007002) were added to the supernatants and rotated at 4 °C overnight. Then the samples were washed 5 times with washing buffer (50 mM HEPES pH7.4, 100 mM NaCl, 0.01% NP-40, 10% glycerol, and 1 mM EDTA) before being boiled for Western blot analysis.

### CUT&Tag and ChIP-seq analysis

CUT&Tag was performed to analyze the genome-wide distribution of Bmi1, RNA Pol II, and histone modifications H2AK119ub, H3K27me3, H3K4me3 in mESCs. Briefly, 10^5^ cells were harvested in NE buffer (20 mM HEPES pH 7.5, 0.5 mM spermidine, 10 mM KCl, 0.1% TritonX-100, 10% Glycerol, and 1 mM PMSF) on ice for 10 min. ConA beads were pre-washed and resuspended in binding buffer (20 mM HEPES pH 7.5, 10 mM KCl, 1 mM CaCl_2_, and 1 mM MnCl_2_). 10 µl of beads were added to each sample and incubated at room temperature for 10 min. Then, the beads were washed with washing buffer (20 mM HEPES pH 7.5, 0.5 mM spermidine, 150 mM NaCl, and 0.1% BSA) and resuspended in blocking buffer (20 mM HEPES pH 7.5, 0.5 mM spermidine, 150 mM NaCl, 0.1% BSA, and 2 mM EDTA) at room temperature for 5 min. Primary antibodies (1:100 dilution) were added and incubated at room temperature for 2 h, followed by washing and incubation with secondary antibodies (1:100 dilution) at room temperature for 30 min. 1.2 µl of PA-Tn5 transposons were added to each sample and incubated at room temperature for 30 min. Beads were resuspended in 30 µl washing buffer with 10 mM MgCl_2_ and incubated at 37 L for 1 h. Reactions were stopped with 5.5 µl stop buffer (2.25 µl of 0.5 M EDTA, 2.75 µl of 10% SDS, and 0.5 µl of 20 mg/ml Proteinase K) and incubated at 55 L for 30 min, and then 70 L for 20 min to inactivate Proteinase K. Tagmented DNA was extracted using 0.9×VAHTS DNA clean beads (VAHTS, Cat. #N411-03).

ChIP-seq was conducted to assess the genome-wide enrichment of histone H2BK120ub. Cells were crosslinked with 1% paraformaldehyde for 10 min, quenched with 125 mM glycine for 5 min at room temperature with rotation, and washed with cold PBS. Cells were then lysed in lysis buffer (10 mM Tris-HCl pH 7.5, 10 mM NaCl, and 0.5% NP-40) on ice for 10 min, and the nuclei were pelleted and resuspended in 500 µl of 1× ChIP-seq buffer (50 mM Tris-HCl pH 8.1, 10 mM EDTA, 100 mM NaCl, 1% Triton X-100, and 0.1% sodium deoxycholate). Chromatin was fragmented by sonication using a Bioruptor (5 cycles of 30 s on/ 30 s off). After centrifugation (15,000 g, 4 °C, 10 min), the supernatant was collected for immunoprecipitation. Chromatin from *Drosophila* S2 cells was spiked in for normalization at 1% of the input. 10 µg of H2BK120ub antibodies (Cell Signaling Technology, Cat. #5546S; Active Motif, Cat. #39623) were added into each sample and rocked at 4 L overnight. Protein A beads were added and incubated for 3 h. After the beads were extensively washed, the bound DNA was eluted in elution buffer (10 mM Tris-HCl pH 8.0, 10 mM EDTA, 150 mM NaCl, 5 mM DTT, and 1% SDS). Crosslinks were reversed by incubation at 65 °C overnight, followed by proteinase K and RNase A treatment. DNA was purified using the MinElute PCR Purification Kit (Qiagen, Cat. #28006) and subjected to library preparation with the VAHTS Universal DNA Library Prep Kit for Illumina V3 (Vazyme, Cat. #ND607-1) following the manufacturer’s instructions.

For CUT&Tag and ChIP-seq data analyses, Trim Galore (version 0.6.7) (https://www.bioinformatics.babraham.ac.uk/projects/trim_galore) with the parameters “-q 20 –paired” was used to remove adaptors and low-quality reads. Trimmed reads were then mapped to the mouse reference genome mm9 using Bowtie2^63^ (version 2.3.5.1). PCR duplicates were removed using GATK4 (version v4.3.0.0) (https://github.com/broadinstitute/gatk) with the parameter “–REMOVE_DUPLICATESLtrue” for ChIP-seq, but not for CUT&Tag data. Genome coverage bigWig files were generated using BEDtools^64^ (version 2.26.0) and bedGraphToBigWig (version 4) (https://www.encodeproject.org/software/bedgraphtobigwig/) with the following parameters “genomecov –scaleFactor 1e^6^ / (the number of reads mapped to the *E.coli* genome (for CUT&Tag) or the *D. melanogaster* genome (for ChIP-seq))”. Specifically, to further eliminate the batch effects between IP and input samples in ChIP-seq analysis, bigWig files of IP samples were further normalized by multiplying the spike-in ratio (*D. melanogaster*/mouse) from input samples. DeepTools^65^ (version 3.5.1) was used to visualize epigenomic signals to draw heat maps using the functions computeMatrix and plotHeatmap. Normalized signals were visualized in Integrative Genomics Viewer (IGV)^66^ (version 2.15.2). A 1 kb sliding window across the whole genome was used to calculate the Pearson product moment correlation between replicate samples.

### RNA-seq analysis

Total RNA was extracted with TRIzol reagent (Sangon Biotech, Cat. #B610409). 500 ng RNA was reversely transcribed into cDNA by Maxima H Minus (Thermo Fisher, Cat. #EP0751). The cDNA was digested using 2 µl of Tn5 transposons in 20 µl 1× DMF buffer at 37 L for 5 min. Reactions were stopped by adding 5.5 µl stop buffer (2.25 µl of 0.5 M EDTA, 2.75 µl of 10% SDS, and 0.5 µl of 20 mg/ml Proteinase K) and incubated at 55 L for 30 min, and then 70 L for 20 min to inactivate Proteinase K. 0.9× of VAHTS DNA clean beads were added to each sample to extract the tagmentated DNA.

RNA-seq raw data were trimmed using Trim Galore as described above. STAR^67^ (version 2.7.6a) was used to map the filtered reads to mouse reference genome mm9. Expression counts were generated using featureCounts^68^ (version 2.0.1) and normalized to reads per kilobase per million mapped reads (RPKM) by R package edgeR^69^ (version 4.2.1). Principal Component Analysis (PCA) was performed by R package gmodels (version 2.19.1) (https://github.com/cran/gmodels) to assess the overall distribution of the samples. The Venn diagram was plotted using the R package VennDiagram (version 1.7.3) (https://github.com/cran/VennDiagram). The gene expression patterns of different groups were visualized by the R package pheatmap (version 1.0.12) (https://github.com/raivokolde/pheatmap). clusterProfiler^70^ (version 4.12.3) was used to perform Gene Ontology (GO) analysis. WordCloud plots were generated by R package wordcloud2 (version 2.19.1) (https://github.com/Lchiffon/wordcloud2). RNA-seq data for mESC cells at different differentiation time points (0, 4, 7, and 14 days). were downloaded from the GEO database GSE271097.

### TT-seq analysis

For TT-seq analysis, mESCs were cultured in 10 cm dishes with 1 mM 4-thiouridine (4-SU, Aladdin, Cat. #T122953) for 15 min. Cells were then collected for RNA extraction using TRIzol reagent. A total of 100 µg of 4-SU-labeled mESC RNA and 1 µg of 4-SU-labeled *Drosophila* S2 RNA were mixed in 100 µl of RNase-free water. 20 µl of 1 M NaOH was added into the mixture and incubated on ice for 20 min, followed by RNA purification using Phenol/Chloroform (pH 6.7). The purified RNA (100 µl) was then mixed with 3 µl of biotin buffer (1 M Tris-HCl pH 7.4 and 0.5 M EDTA) and 50 µl of 0.1 mg/ml Biotin MTSEA (Biotium, Cat. #90064-90066-1). After another round of phenol/chloroform (pH 6.7) purification, RNA was reconstituted in 50 µl of RNase-free water, incubated at 65 °C for 10 min, and rapidly cooled on ice for 5 min. Pre-washed streptavidin beads (Smart-lifesciences, Cat. #SM007002) were then added to the supernatant and rotated at room temperature for 1 h in the dark. The beads were washed five times with high-salt washing buffer (100 mM Tris-HCl pH 7.4, 1 M NaCl, 0.1% Tween-20, and 10 mM EDTA) before RNA was eluted twice using 100 µl of 100 mM DTT (Sangon Biotech, Cat. #A620058). The eluted RNA was subsequently purified via ethanol precipitation and subjected to library preparation using an RNA-seq library prep kit (VAHTS, Cat. #NR604), following the manufacturer’s instructions.

TT-seq raw data were trimmed using Trim Galore as described above. Trimmed reads were uniquely mapped to the mouse reference genome (mm9) using Bowtie2^63^ (version 2.3.5.1). Genome coverage bigWig files were generated by bamCoverage (deeptools 3.5.1) with the parameter ‘--scaleFactor 1e^6^ / (the number of reads mapped to the *D. melanogaster* genome)’ and the parameter ‘--filterRNAstrand’ to retain specific strand information. HOMER^71^ (v4.11.1) was used to generate the expression profiles using the annotatePeaks.pl function with the parameter “-size 2000 -hist 10 - pc 5”.

### Calculation of RNA Pol II pausing index

The promoter-proximal bin was defined as a fixed window from -500 bp to +100 bp around the TSS. The transcribed region (gene body) bin was defined from +500 bp to the TES. The index for each gene was calculated as the ratio of Pol II density in the promoter-proximal bin to the Pol II density in the transcribed region bin.

### Antibodies

Rabbit polyclonal anti-Histone H3 (Abcam, Cat. #ab1791); Rabbit polyclonal anti-FLAG Tag (Sangon Biotech, Cat. #D110005); Rabbit polyclonal anti-Histone H3K4me3 (Abcam, Cat. #ab8580); Rabbit monoclonal anti-Histone H3K27me3 (Cell Signaling Technology, Cat. #9733); Rabbit monoclonal anti-H2AK119ub (Cell Signaling Technology, Cat. #8240); Rabbit monoclonal anti-H2BK120ub (Cell Signaling Technology, Cat. #5546S); Rabbit monoclonal anti-H2BK120ub (Active Motif, Cat. #39623); Mouse monoclonal anti-POLR2 (EMD Millipore, Cat. #05623); Rabbit monoclonal anti-Bmi1 (Cell Signaling Technology, Cat. #5856); Rabbit monoclonal anti-RNF20 (Cell Signaling Technology, Cat. #11974). Peroxidase AffiniPure Goat anti-Rabbit IgG (H + L) (Jackson ImmunoResearch Laboratories, Cat. #111-035-003); Peroxidase AffiniPure Goat anti-Mouse IgG (H + L) (Jackson ImmunoResearch Laboratories, Cat. #115-035-003); Alexa Fluor® 488 AffiniPure Alpaca Anti-Rabbit IgG (H + L) (Jackson ImmunoResearch Laboratories, Cat. #611-035-215).

## Data availability

The cryo-EM maps and coordinates have been deposited into the Electron Microscopy Data Bank (EMDB) and the Protein Data Bank (PDB) as follows: cPRC1 E3 (RNF2-BMI1) : E2∼ub : NCP^H2BK120ub^ complex (EMD-XXXXX; PDB XXXX); ncPRC1.1 E3 (RNF2-PCGF1) : E2∼ub : NCP^H2BK120ub^ complex (EMD-XXXXX; PDB XXXX); ncPRC1.1-NCP^H2BK120ub&H2AK119ub^ complex (EMD-XXXXX; PDB XXXX); ncPRC1.6-NCP^H2BK120ub&H2AK119ub^ complex (EMD-XXXXX; PDB XXXX); ncPRC1.4-NCP^H2BK120ub&H2AK119ub^ complex_2 RNF2-BMI1 (EMD-XXXXX; PDB XXXX); ncPRC1.4-NCP^H2BK120ub&H2AK119ub^ complex_1 RNF2-BMI1 & 1 RYBP (EMD-XXXXX; PDB XXXX); ncPRC1.4-NCP^H2AK119ub^ complex_1 RNF2-BMI1 (EMD-XXXXX; PDB XXXX); ncPRC1.4-NCP^H2AK119ub^ complex_1 RNF2-BMI1 & 1 RYBP (EMD-XXXXX; PDB XXXX); ncPRC1.4-NCP^H2AK119ub^ complex_1 RYBP (EMD-XXXXX; PDB XXXX); ncPRC1.4-NCP^H2AK119ub^ complex_2 RYBP (EMDrXXXXX; PDB XXXX). The raw and processed sequencing data generated in this study have been deposited in the NCBI Gene Expression Omnibus (GEO) database under accession GSEXXXXXX, GSEXXXXXX, GSEXXXXXX, and GSEXXXXXX.

## References

1. Kim, J. J. & Kingston, R. E. Context-specific Polycomb mechanisms in development. Nat. Rev. Genet. 23, 680–695 (2022).

2. Piunti, A. & Shilatifard, A. The roles of Polycomb repressive complexes in mammalian development and cancer. Nat. Rev. Mol. Cell Biol. 22, 326–345 (2021).

3. Blackledge, N. P. & Klose, R. J. The molecular principles of gene regulation by Polycomb repressive complexes. Nat. Rev. Mol. Cell Biol. 22, 815–833 (2021).

4. Schuettengruber, B., Bourbon, H. M., Di Croce, L. & Cavalli, G. Genome regulation by Polycomb and Trithorax: 70 years and counting. Cell 171, 34–57 (2017).

5. Chan, H. L. & Morey, L. Emerging roles for Polycomb-group roteins in stem cells and cancer. Trends Biochem. Sci. 44, 688–700 (2019).

6. Wang, H., et al. Role of histone H2A ubiquitination in Polycomb silencing. Nature 431, 862–868 (2004).

7. Napoles, M. De et al. Polycomb group proteins Ring1A / B link ubiquitylation of histone H2A to heritable gene silencing and X inactivation. Dev. Cell 7, 663–676 (2004).

8. Müller, J. et al. Histone methyltransferase activity of a Drosophila Polycomb group repressor complex. Cell 111, 197–208 (2002).

9. Cao, R., Wang, L., Wang, H. & Xia, L. Role of histone H3 lysine 27 methylation in Polycomb-group silencing. Science 298, 1039–1043 (2002).

10. Blackledge, N. P. et al. Variant PRC1 complex-dependent H2A ubiquitylation drives PRC2 recruitment and polycomb domain formation. Cell 157, 1445–1459 (2014).

11. Gao, Z. et al. PCGF homologs, CBX proteins, and RYBP define functionally distinct PRC1 family complexes. Mol. Cell 45, 344–356 (2012).

12. Tavares, L. et al. RYBP-PRC1 complexes mediate H2A ubiquitylation at polycomb target sites independently of PRC2 and H3K27me3. Cell 148, 664–678 (2012).

13. Margueron, R. et al. Role of the polycomb protein EED in the propagation of repressive histone marks. Nature 461, (2009).

14. Sauer, P. V. et al. Activation of automethylated PRC2 by dimerization on chromatin. Mol. Cell 84, 3885–3898 (2024).

15. Kasinath, V. et al. JARID2 and AEBP2 regulate PRC2 in the presence of H2AK119ub1 and other histone modifications. Science 371, (2021).

16. Ciapponi, M., Karlukova, E., Schkölziger, S., Benda, C. & Müller, J. Structural basis of the histone ubiquitination read-write mechanism of RYBP-PRC1. Nat. Struct. Mol. Biol. 31, 1023–1027 (2024).

17. López, V. G. et al. Read-write mechanisms of H2A ubiquitination by Polycomb repressive complex 1. Nature 636, 755–761 (2024).

18. Shilatifard, A. The COMPASS family of histone H3K4 methylases: mechanisms of regulation in development and disease pathogenesis. Annu. Rev. Biochem. 81, 65–95 (2012).

19. Function, G. & Kennison, J. A. The Polycomb and Trithorax group proteins of Drosophila: trans-regulators of homeotic gene function. Annu. Rev. Genet. 29, 289–303 (1995).

20. Piunti, A. & Shilatifard, A. Epigenetic balance of gene expression by polycomb and compass families. Science 352, (2016).

21. Sun, Z.-W. & Allis, C. D. Ubiquitination of histone H2B regulates H3 methylation and gene silencing in yeast. Nature 418, 104–108 (2002).

22. Kim, J. et al. RAD6-mediated transcription-coupled H2B ubiquitylation directly stimulates H3K4 methylation in human cells. Cell 137, 459–471 (2009).

23. Xue, H. et al. Structural basis of nucleosome recognition and modification by MLL methyltransferases. Nature 573, 445–449 (2019).

24. Hsu, P. L. et al. Structural basis of H2B ubiquitination-dependent H3K4 methylation by COMPASS. Mol. Cell 76, 712–723 (2019).

25. Macrae, T. A., Fothergill-Robinson, J. & Ramalho-Santos, M. Regulation, functions and transmission of bivalent chromatin during mammalian development. Nat. Rev. Mol. Cell Biol. 24, 6–26 (2023).

26. Bernstein, B. E. et al. A bivalent chromatin structure marks key developmental genes in embryonic stem cells. Cell 125, 315–326 (2006).

27. Azuara, V. et al. Chromatin signatures of pluripotent cell lines. Nat. Cell Biol. 8, (2006).

28. Boyer, L. A. et al. Polycomb complexes repress developmental regulators in murine embryonic stem cells. Nature 441, 349–353 (2006).

29. Kaustov, L. et al. Recognition and specificity determinants of the human Cbx chromodomains. J. Biol. Chem. 286, 521–529 (2011).

30. Kim, C. A., Gingery, M., Pilpa, R. M. & Bowie, J. U. The SAM domain of polyhomeotic forms a helical polymer. Nat. Struct. Biol. 9, 453–457 (2002).

31. Isono, K. et al. SAM domain polymerization links subnuclear clustering of PRC1 to gene silencing. Dev. Cell 26, 565–577 (2013).

32. Seif, E. et al. Phase separation by the polyhomeotic sterile alpha motif compartmentalizes Polycomb Group proteins and enhances their activity. Nat. Commun. 11, 1–19 (2020).

33. Boettiger, A. N. et al. Super-resolution imaging reveals distinct chromatin folding for different epigenetic states. Nature 529, 418–422 (2016).

34. Kingston, R. E., Francis, N. J., Sadreyev, R. I. & Zhuang, X. Chromatin topology is coupled to Polycomb group protein subnuclear organization. Nat. Commun. 7, 1–13 (2016).

35. Kundu, S. et al. Polycomb repressive complex 1 generates discrete compacted domains that change during differentiation. Mol. Cell 65, 432–446 (2017).

36. Zhao, J. et al. RYBP/YAF2-PRC1 complexes and histone H1-dependent chromatin compaction mediate propagation of H2AK119ub1 during cell division. Nat. Cell Biol. 22, 439–452 (2020).

37. McGinty, R. K., Henrici, R. C. & Tan, S. Crystal structure of the PRC1 ubiquitylation module bound to the nucleosome. Nature 514, 591–596 (2014).

38. Yao, S., Jeon, Y., Kesner, B. & Lee, J. T. Xist RNA binds select autosomal genes and depends on Repeat B to regulate their expression. Elife 1–48 (2024).

39. McGinty, R. K., Kim, J., Chatterjee, C., Roeder, R. G. & Muir, T. W. Chemically ubiquitylated histone H2B stimulates hDot1L-mediated intranucleosomal methylation. Nature 453, 812–6 (2008).

40. Worden, E. J., Hoffmann, N. A., Hicks, C. W. & Wolberger, C. Mechanism of cross-talk between H2B ubiquitination and H3 methylation by Dot1L. Cell 176, 1490–1501.e12 (2019).

41. Hicks, C. W. et al. Ubiquitinated histone H2B as gatekeeper of the nucleosome acidic patch. Nucleic Acids Res. 52, 9978–9995 (2024).

42. Morey, L. et al. Polycomb regulates mesoderm cell fate-specification in embryonic stem cells through activation and repression mechanisms. Cell Stem Cell 17, 300–315 (2015).

43. Loubiere, V. et al. Coordinate redeployment of PRC1 proteins suppresses tumor formation during Drosophila development. Nat. Genet. 48, 1436–1442 (2016).

44. Pherson, M. et al. Polycomb repressive complex 1 modifies transcription of active genes. Sci. Adv. 3, 1–17 (2017).

45. Cohen, I. et al. PRC1 fine-tunes gene repression and activation to safeguard skin development and stem cell specification. Cell Stem Cell 22, 726–739 (2018).

46. Chan, H. L. et al. Polycomb complexes associate with enhancers and promote oncogenic transcriptional programs in cancer through multiple mechanisms. Nat. Commun. 9, 1–16 (2018).

47. Loubiere, V., Papadopoulos, G. L., Szabo, Q., Martinez, A. M. & Cavalli, G. Widespread activation of developmental gene expression characterized by PRC1-dependent chromatin looping. Sci. Adv. 6, 1–12 (2020).

48. Dyer, P. N. et al. Reconstitution of nucleosome core particles from recombinant histones and DNA. Methods Enzymol. 375, 23–44 (2004).

49. Morgan, M. T. et al. Structural basis for histone H2B deubiquitination by the SAGA DUB module. Science 351, 725–728 (2016).

50. Kastner, B. et al. GraFix: sample preparation for single-particle electron cryomicroscopy. Nat. Methods 5, 53–55 (2008).

51. Zheng, S. Q. et al. MotionCor2: anisotropic correction of beam-induced motion for improved cryo-electron microscopy. Nat. Methods 14, 331–332 (2017).

52. Grant, T. & Grigorieff, N. Measuring the optimal exposure for single particle cryo-EM using a 2.6 Å reconstruction of rotavirus VP6. Elife 4, 1–19 (2015).

53. Rohou, A. & Grigorieff, N. CTFFIND4: Fast and accurate defocus estimation from electron micrographs. J. Struct. Biol. 192, 216–221 (2015).

54. Bepler, T. et al. Positive-unlabeled convolutional neural networks for particle picking in cryo-electron micrographs. Nat. Methods 16, 1153–1160 (2019).

55. Kimanius, D., Dong, L., Sharov, G., Nakane, T. & Scheres, S. H. W. New tools for automated cryo-EM single-particle analysis in RELION-4.0. Biochem. J. 478, 4169–4185 (2021).

56. Bai, X. C., Rajendra, E., Yang, G., Shi, Y. & Scheres, S. H. W. Sampling the conformational space of the catalytic subunit of human g-secretase. Elife 4, 1–19 (2015).

57. Scheres, S. H. W. & Chen, S. Prevention of overfitting in cryo-EM structure determination. Nat. Methods 9, 853–854 (2012).

58. Kucukelbir, A., Sigworth, F. J. & Tagare, H. D. Quantifying the local resolution of cryo-EM density maps. Nat. Methods 11, 63–65 (2014).

59. Pettersen, E. F. et al. UCSF Chimera - A visualization system for exploratory research and analysis. J. Comput. Chem. 25, 1605–1612 (2004).

60. Emsley, P., Lohkamp, B., Scott, W. G. & Cowtan, K. Features and development of Coot. Acta Crystallogr. Sect. D Biol. Crystallogr. 66, 486–501 (2010).

61. Adams, P. D. et al. PHENIX: A comprehensive Python-based system for macromolecular structure solution. Acta Crystallogr. Sect. D Biol. Crystallogr. 66, 213–221 (2010).

62. Wan, X. et al. MAT2B regulates the protein level of MAT2A to preserve RNA N6-methyladenosine. Cell Death Dis. 15, 714 (2024).

63. Langmead, B. & Salzberg, S. L. Fast gapped-read alignment with Bowtie 2. Nat. Methods 9, 357–359 (2012).

64. Quinlan, A. R. BEDTools: The Swiss-Army tool for genome feature analysis. Current Protocols in Bioinformatics vol. 47 (2014).

65. Ramírez, F., Dündar, F., Diehl, S., Grüning, B. A. & Manke, T. DeepTools: a flexible platform for exploring deep-sequencing data. Nucleic Acids Res. 42, 187–191 (2014).

66. Thorvaldsdóttir, H., Robinson, J. T. & Mesirov, J. P. Integrative Genomics Viewer (IGV): High-performance genomics data visualization and exploration. Brief. Bioinform. 14, 178–192 (2013).

67. Dobin, A. et al. STAR: Ultrafast universal RNA-seq aligner. Bioinformatics 29, 15–21 (2013).

68. Liao, Y., Smyth, G. K. & Shi, W. FeatureCounts: An efficient general purpose program for assigning sequence reads to genomic features. Bioinformatics 30, 923–930 (2014).

69. Robinson, M. D., McCarthy, D. J. & Smyth, G. K. edgeR: A Bioconductor package for differential expression analysis of digital gene expression data. Bioinformatics 26, 139–140 (2009).

70. Yu, G., Wang, L. G., Han, Y. & He, Q. Y. ClusterProfiler: An R package for comparing biological themes among gene clusters. Omi. A J. Integr. Biol. 16, 284–287 (2012).

71. Heinz, S. et al. Simple combinations of lineage-determining transcription factors prime cis-regulatory elements required for macrophage and B cell identities. Mol. Cell 38, 576–589 (2010).

