## Extended data figures for "Histone H2BK120 ubiquitination modulates PRC1 activity and H2AK119ub deposition on nucleosomes"

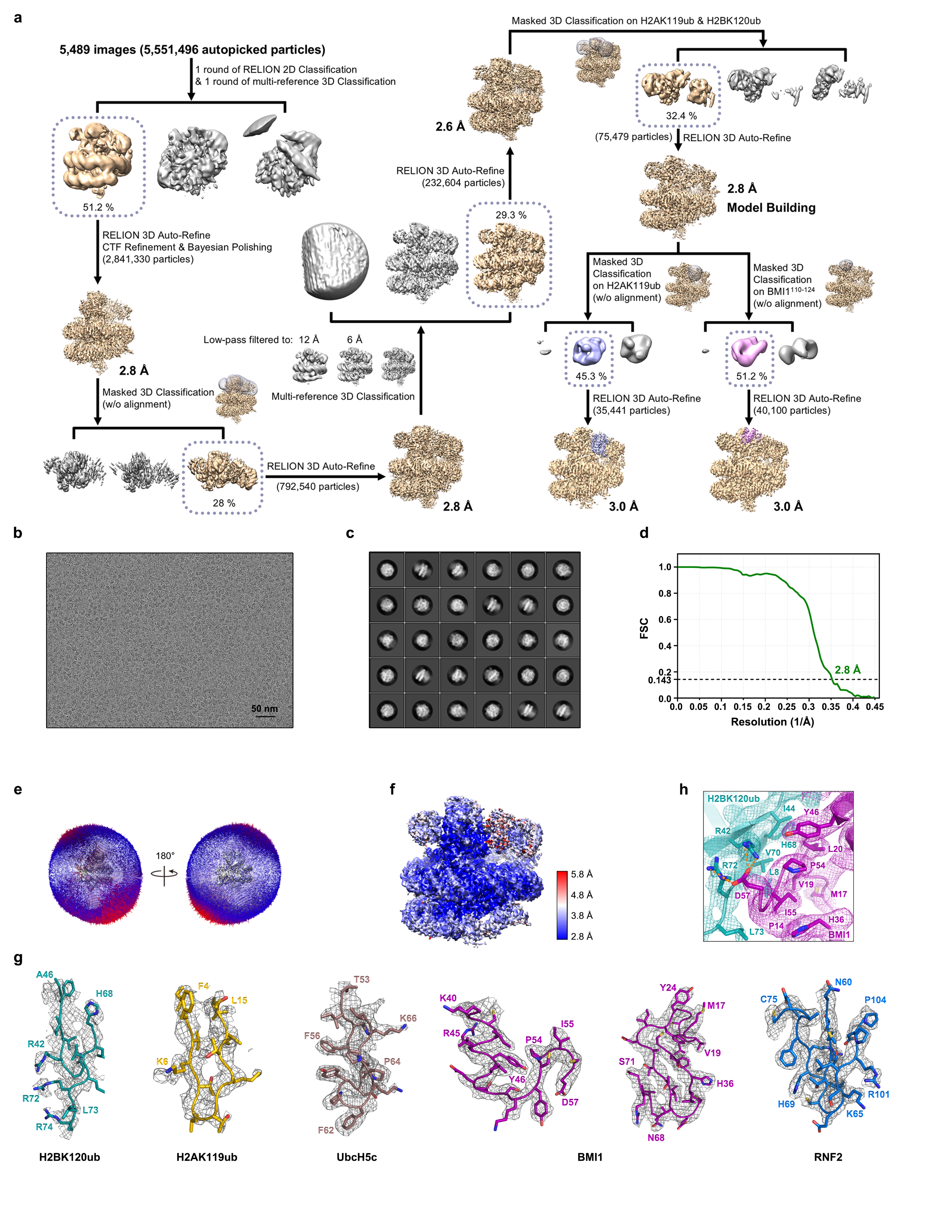

**Extended Data Fig. 1 |** **Structural characterization of human cPRC1–E2~ub-NCP^H2BK120ub^ complex. a,** Flow chart of cryo-EM data processing of the cPRC1-E2~ub-NCP^H2BK120ub^ dataset. **b**, Representative micrograph of the cryo-EM dataset of the cPRC1-E2~ub-NCP^H2BK120ub^ complex. **c**, Representative 2D class averages of cryo-EM particles of the cPRC1-E2~ub-NCP^H2BK120ub^ complex. **d**, The gold-standard FSC curve calculated between two halves of the cPRC1-E2~ub-NCP^H2BK120ub^ dataset. **e**, Angular distribution of particle projections of the cPRC1-E2~ub-NCP^H2BK120ub^ reconstruction. **f**, Local-resolution estimates of the cPRC1-E2~ub-NCP^H2BK120ub^ complex. **g**, Representative electron microscopy (EM) density maps of the cPRC1-E2~ub-NCP^H2BK120ub^ complex structure. **h**, EM densities of residues at the H2BK120ub-BMI1 interface, related to Fig. 1e.

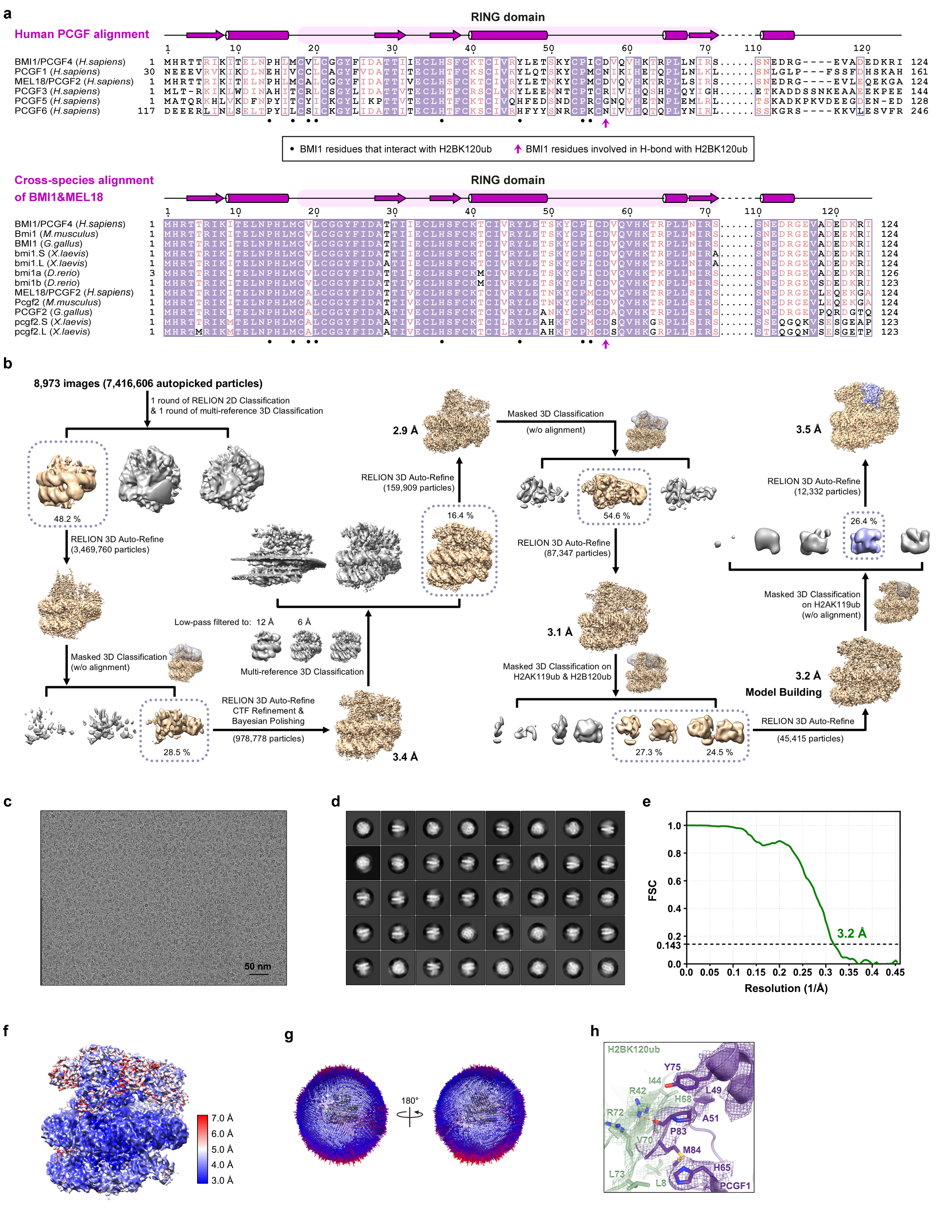

**Extended Data Fig. 2 |** **Structural characterization of human ncPRC1.1–E2~ub-NCP^H2BK120ub^ complex. a**, Sequence alignment of the RING domains of human PCGFs (top) and cross-species alignment of the RING domains of BMI1 and MEL18 (bottom), with identical residues highlighted in purple and conserved residues colored in pink. The secondary structure of the RING domain is shown above the sequence alignment. **b**, Flow chart of cryo-EM data processing of the ncPRC1.1-E2~ub-NCP^H2BK120ub^ dataset. **c**, Representative micrograph of the cryo-EM dataset of the ncPRC1.1-E2~ub-NCP^H2BK120ub^ complex. **d**, Representative 2D class averages of cryo-EM particles of the ncPRC1.1-E2~ub-NCP^H2BK120ub^ complex. **e**, The gold-standard FSC curve calculated between two halves of the ncPRC1.1-E2~ub-NCP^H2BK120ub^ dataset. **f**, Local-resolution estimates of the ncPRC1.1-E2~ub-NCP^H2BK120ub^ complex. **g**, Angular distribution of particle projections of the ncPRC1.1-E2~ub-NCP^H2BK120ub^ reconstruction. **h**, EM densities of residues at the H2BK120ub-PCGF1 interface, related to Fig. 2e.

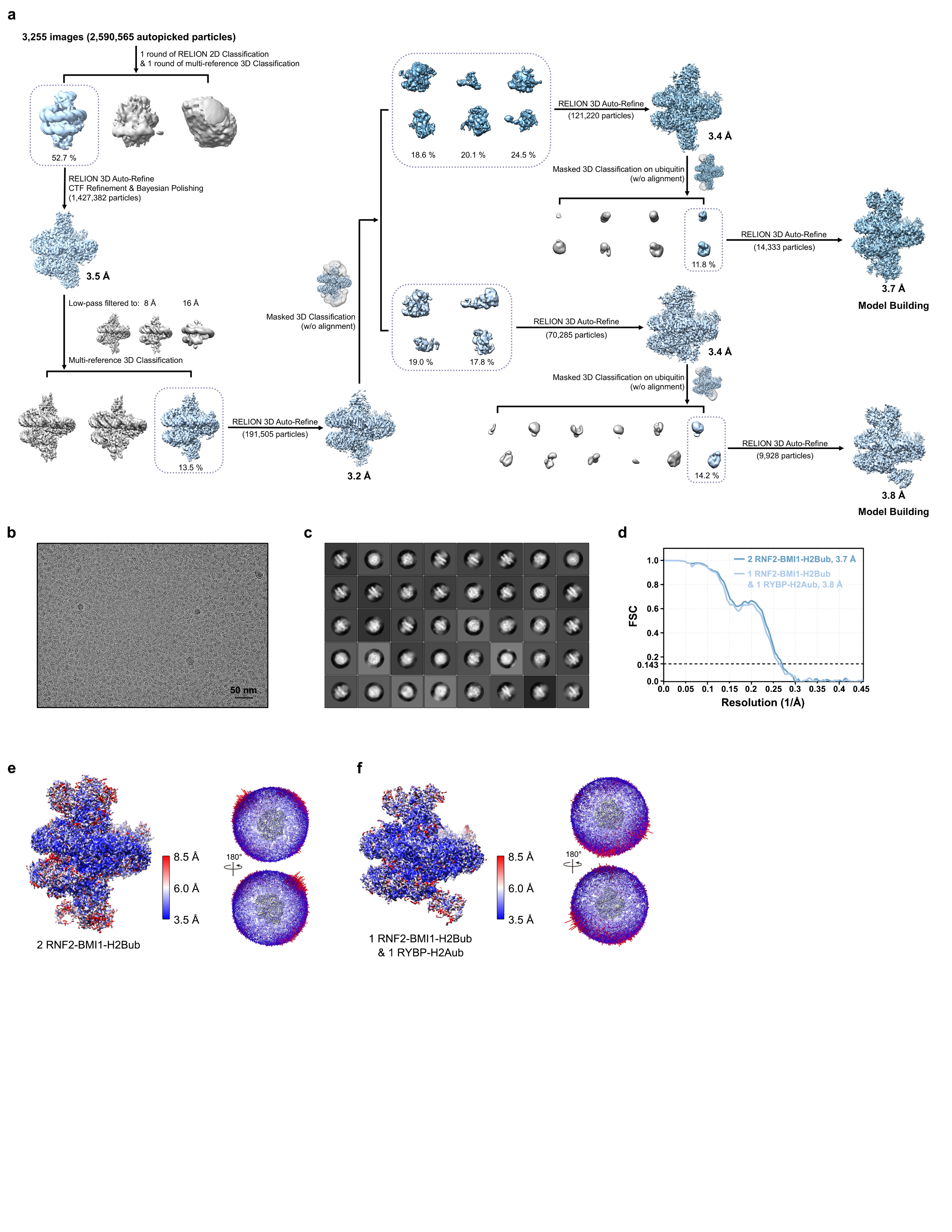

**Extended Data Fig. 3 |** **Structural characterization of human ncPRC1.4-NCP^H2BK120ub&H2AK119ub^ complex. a,** Flow chart of cryo-EM data processing of the ncPRC1.4-NCP^H2BK120ub&H2AK119ub^ complex dataset. **b**, Representative micrograph of the cryo-EM dataset of the ncPRC1.4-NCP^H2BK120ub&H2AK119ub^ complex. **c**, Representative 2D class averages of cryo-EM particles of the ncPRC1.4-NCP^H2BK120ub&H2AK119ub^ complex. **d**, The gold-standard FSC curve calculated between two halves of the ncPRC1.4-NCP^H2BK120ub&H2AK119ub^ complex dataset. **e**, Local-resolution estimates (left) and angular distribution (right) of particle projections of the ncPRC1.4-NCP^H2BK120ub&H2AK119ub^ complex which contains two RNF2-BMI1 modules binding to opposite sides of a single NCP^H2BK120ub&H2AK119ub^ (denoted as 2 RNF2-BMI1-H2Bub). **f**, Local-resolution estimates (left) and angular distribution (right) of particle projections of the ncPRC1.4-NCP^H2BK120ub&H2AK119ub^ complex which contains one RNF2-BMI1 module and one RYBP subunit bind to opposite sides of a single NCP^H2BK120ub&H2AK119ub^ (denoted as 1 RNF2-BMI1-H2Bub & 1 RYBP-H2Aub).

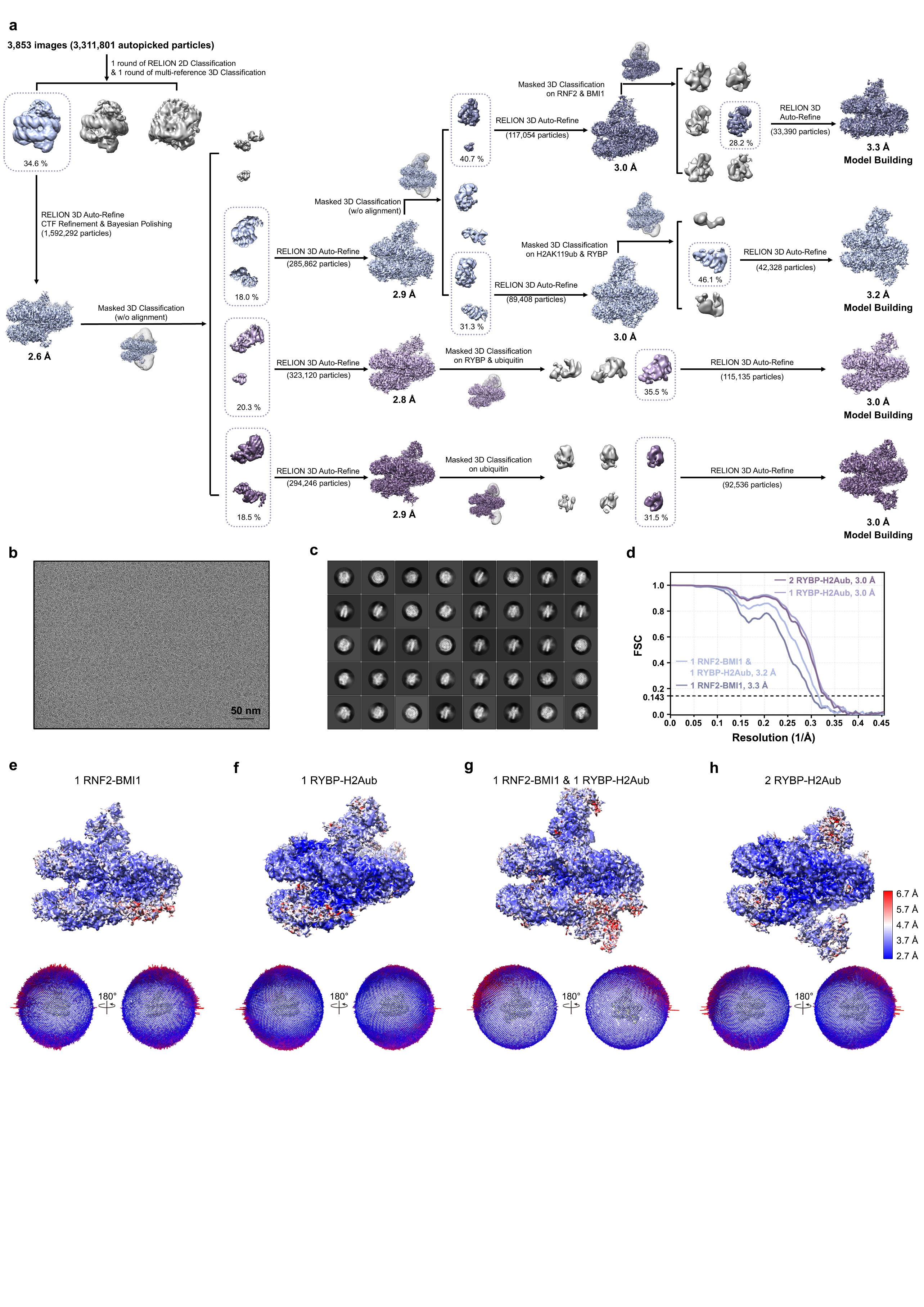

**Extended Data Fig. 4 |** **Structural characterization of human ncPRC1.4-NCP^H2AK119ub^ complex. a**, Flow chart of cryo-EM data processing of the ncPRC1.4-NCP^H2AK119ub^ complex dataset. **b**, Representative micrograph of the cryo-EM dataset of the ncPRC1.4-NCP^H2AK119ub^ complex. **c**, Representative 2D class averages of cryo-EM particles of the ncPRC1.4-NCP^H2AK119ub^ complex. **d,** The gold-standard FSC curve calculated between two halves of the ncPRC1.4-NCP^H2AK119ub^ complex dataset. **e**, Local-resolution estimates (top) and angular distribution (bottom) of particle projections of the ncPRC1.4-NCP^H2AK119ub^ complex which contains one RNF2-BMI1 module binds to one side of NCP^H2AK119ub^ (1 RNF2-BMI1). **f**, Local-resolution estimates (top) and angular distribution (bottom) of particle projections of the ncPRC1.4-NCP^H2AK119ub^ complex which contains one RYBP subunit binds to one side of NCP^H2AK119ub^ (1 RYBP-H2Aub). **g**, Local-resolution estimates (top) and angular distribution (bottom) of particle projections of the ncPRC1.4-NCP^H2AK119ub^ complex which contains one RNF2-BMI1 module and one RYBP subunit bind to opposite sides of a single NCP^H2AK119ub^ (1 RNF2-BMI1 & 1 RYBP-H2Aub). **h**, Local-resolution estimates (top) and angular distribution (bottom) of particle projections of the ncPRC1.4-NCP^H2AK119ub^ complex which contains two RYBP subunits bind to opposite sides of a single NCP^H2AK119ub^ (2 RYBP-H2Aub).

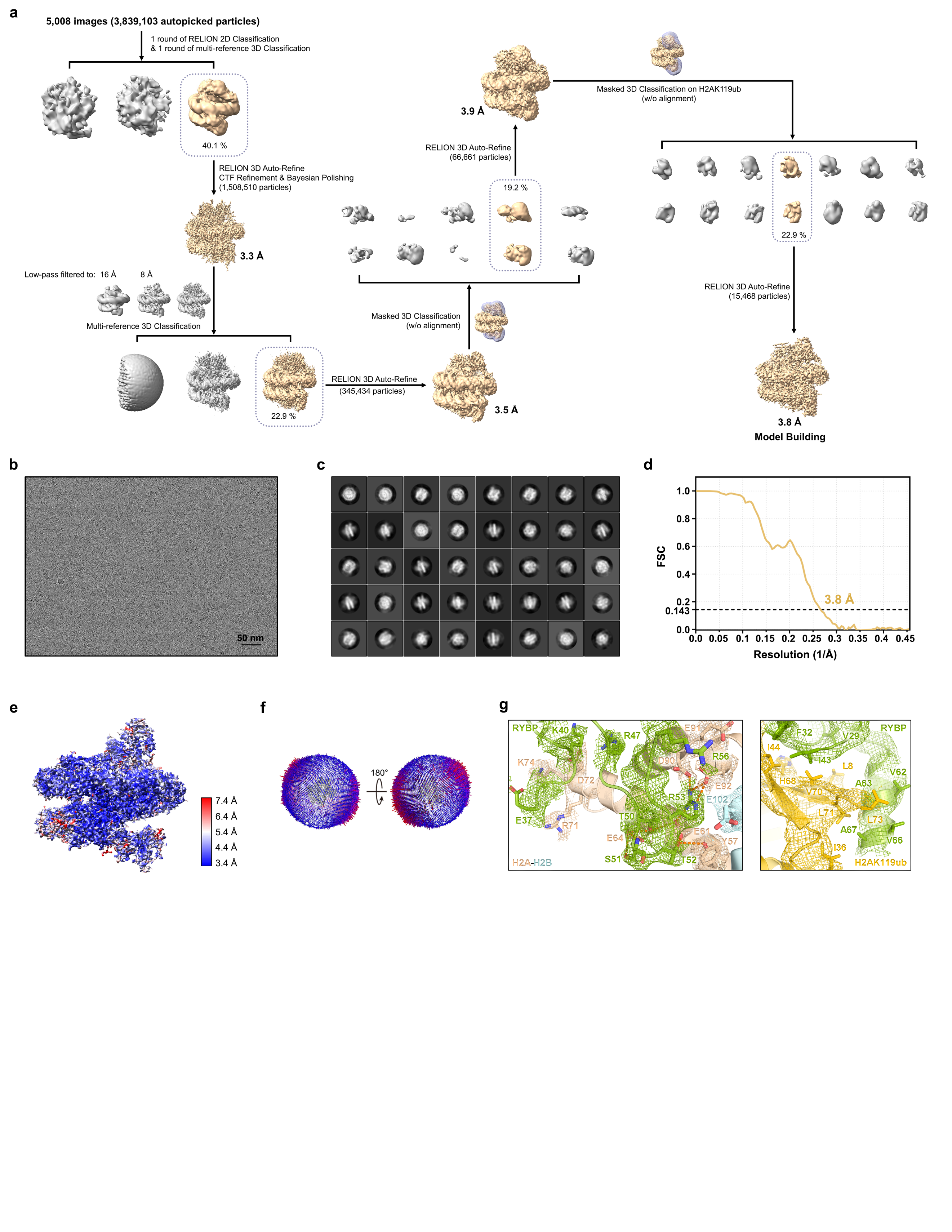

**Extended Data Fig. 5 |** **Structural characterization of human ncPRC1.1-NCP^H2BK120ub&H2AK119ub^ complex. a,** Flow chart of cryo-EM data processing of human ncPRC1.1-NCP^H2AK119ub&H2BK120ub^ dataset. **b**, Representative micrograph of the cryo-EM dataset of the ncPRC1.1-NCP^H2AK119ub&H2BK120ub^ complex. **c**, Representative 2D class averages of cryo-EM particles of the ncPRC1.1-NCP^H2AK119ub&H2BK120ub^ complex. **d**, The gold-standard FSC curve calculated between two halves of the ncPRC1.1-NCP^H2BK120ub&H2AK119ub^ dataset. **e**, Local-resolution estimates of the ncPRC1.1-NCP^H2BK120ub&H2AK119ub^ complex. **f**, Angular distribution of particle projections of the ncPRC1.1-NCP^H2BK120ub&H2AK119ub^ reconstruction. **g**, EM density maps of residues at the RYBP-nucleosome interface (left) and the RYBP-H2AK119ub interface (right), related to Fig. 3b.

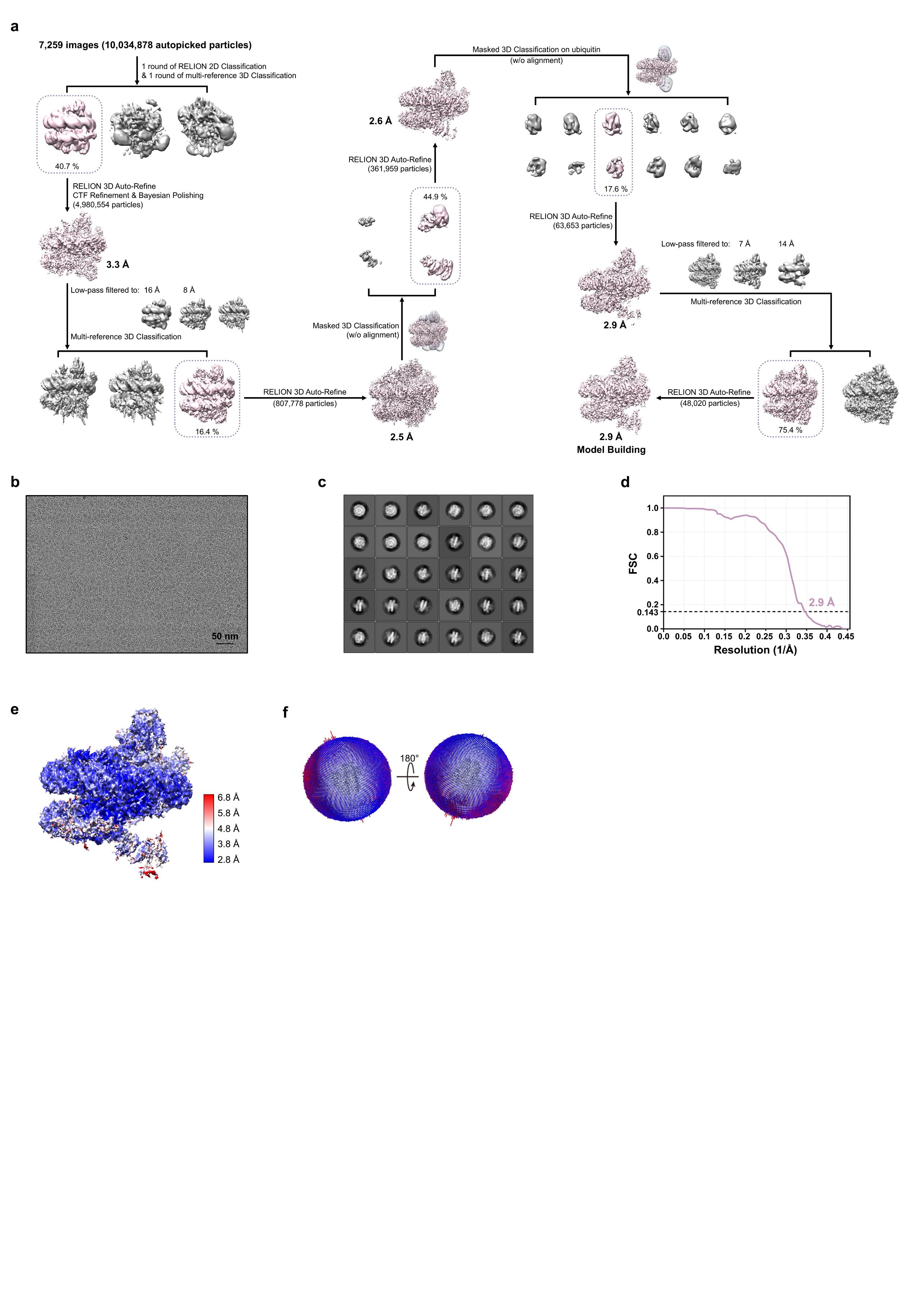

**Extended Data Fig. 6 |** **Structural characterization of human ncPRC1.6-NCP^H2BK120ub&H2AK119ub^ complex. a,** Flow chart of cryo-EM data processing of human ncPRC1.6-NCP^H2AK119ub&H2BK120ub^ dataset. **b**, Representative micrograph of the cryo-EM dataset of the ncPRC1.6-NCP^H2AK119ub&H2BK120ub^ complex. **c**, Representative 2D class averages of cryo-EM particles of the ncPRC1.6-NCP^H2AK119ub&H2BK120ub^ complex. **d**, The gold-standard FSC curve calculated between two halves of the ncPRC1.6-NCP^H2BK120ub&H2AK119ub^ dataset. **e**, Local-resolution estimates of the ncPRC1.6-NCP^H2BK120ub&H2AK119ub^ complex. **f**, Angular distribution of particle projections of the ncPRC1.6-NCP^H2BK120ub&H2AK119ub^ reconstruction.

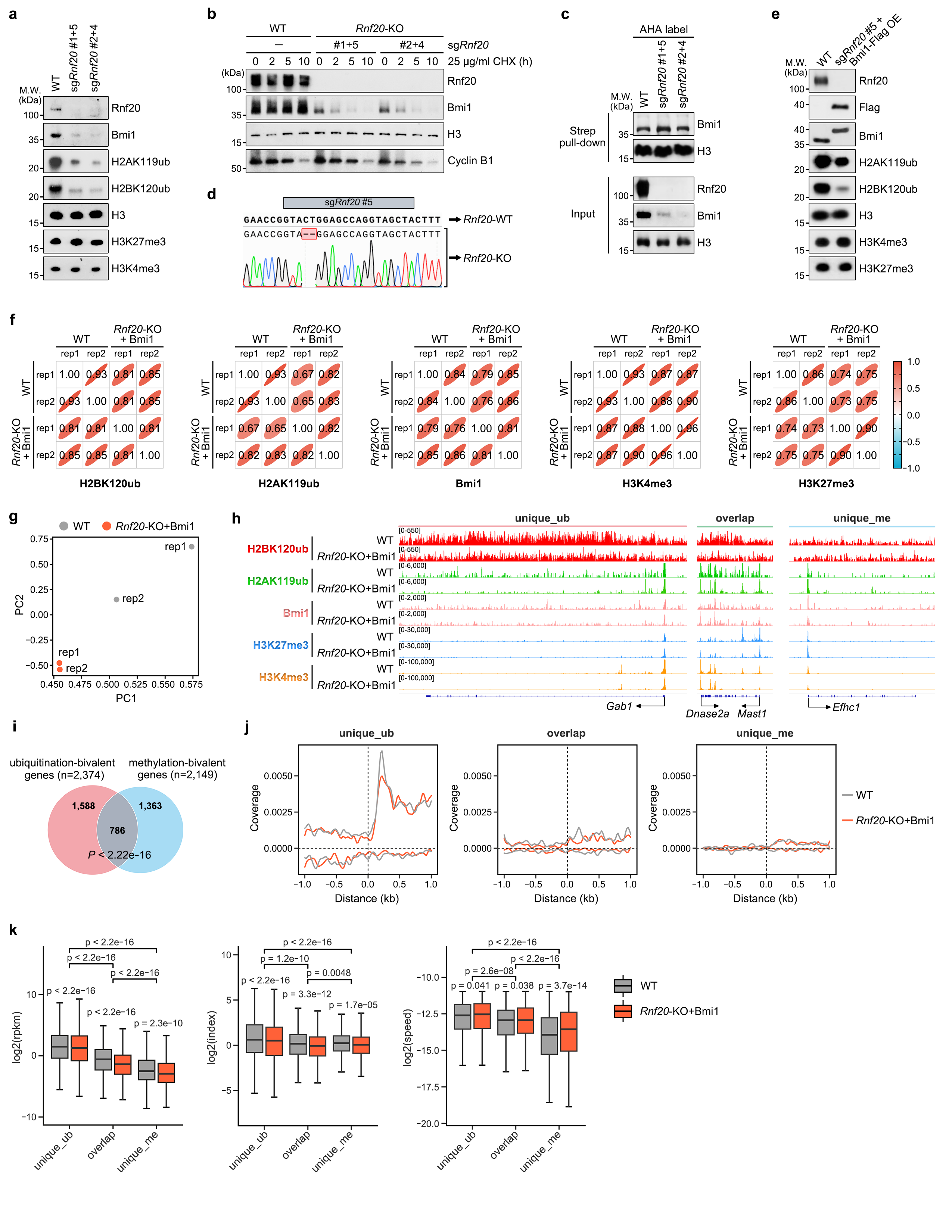

**Extended Data Fig. 7 |** **Genome-wide distribution and gene expression analysis of H2BK120ub and H2AK119ub colocalization in mESCs. a**, Western blot showing that deletion of *Rnf20* decreased the levels of Bmi1, H2AK119ub, and H2BK120ub, while H3K27me3 and H3K4me3 levels remained unchanged. Four independent sgRNAs were used to knockout *Rnf20*. The assay was repeated three times with similar results. **b**, Bmi1 degrades more slowly in WT mESCs compared to two *Rnf20*-KO mESC lines. Cells were collected at 0, 2, 5, and 10 h after 25 μg/mL cycloheximide treatment. CyclinB1 was probed as a positive control. Western blot was conducted using the indicated antibodies. The assay was repeated three times with similar results. **c,** Western blot showing that the translation efficiency of Bmi1 was not altered after the deletion of *Rnf20* in mESCs. Four independent sgRNAs were used to knockout *Rnf20*. The assay was repeated three times with similar results. **d**, Sanger sequencing result of *Rnf20*-KO mESCs. *sgRNF20#5* was shown as a representative example. KO was generated through a 2-base pair deletion. **e,** Western blot showing that re-expression of BmiI1 in *Rnf20-*KO cell line resulted in decreased levels of H2A119ub and H2B120ub, while H3K27me3 and H3K4me3 levels remained unchanged. *Rnf20*-KO cell line was generated by sgRNA through a 2-base pair deletion. The assay was repeated twice with similar results. **f**, Pearson correlations among the ChIP-Seq data of H2BK120ub and the CUT&Tag data of H2AK119ub, Bmi1, H3K4me3, and H3K27me3 from WT and *Rnf20-*KO+Flag-Bmi1 mESCs. The terms “rep1” and “rep2” refer to the technical replicates used in this analysis. **g**, Principal component analysis (PCA) plot of gene expression data from WT and *Rnf20*-KO+Flag-Bmi1 mESCs. Two replicates of each cell line are presented. **h**, Integrative Genomics Viewer (IGV) tracks presenting the enrichment of H2BK120ub, H2AK119ub, Bmi1, H3K27me3 and H3K4me3 at representative loci of unique_ub, overlap, and unique_me categories within WT and *Rnf20-*KO+Flag-Bmi1 mESCs. **i**, Venn diagram illustrating the overlap of di-ubiquitination genes and di-methylation genes in WT mESCs. P value was determined by hypergeometric test. **j**, Profiles showing the average normalized strand-specific coverage around the TSS in different gene groups from WT and *Rnf20*-KO+Flag-Bmi1 mESCs. **k**, Box plots showing the expression of different gene groups (left), the Pol II pausing index (medium), and mRNA synthesis speed (right) of the unique_ub, overlap, and unique_me categories within WT and *Rnf20*-KO+Flag-Bmi1 mESCs. P values within groups were calculated by Student’s t-test (two-sided, paired), and P values between groups were calculated by Student’s t-test (two-sided). The boxes were drawn from lower quartile (Q1) to upper quartile (Q3) with the middle line denoting the median, and whiskers with maximum 1.5 IQR.
